# Human pluripotent stem cell-derived atrioventricular node-like pacemaker cells exhibit biological conduction bridge properties in vitro and in vivo

**DOI:** 10.1101/2025.09.04.674322

**Authors:** Michelle Lohbihler, Amos A. Lim, Stéphane Massé, Maggie Kwan, Omar Mourad, Olya Mastikhina, Brandon M. Murareanu, Malak Elbatarny, Renu Sarao, Beiping Qiang, Wahiba Dhahri, Matthew L. Chang, Alice L.Y. Xu, Amine Mazine, Shahryar Khattak, Sara S. Nunes, Kumaraswamy Nanthakumar, Michael A. Laflamme, Stephanie Protze

**Affiliations:** McEwen Stem Cell Institute, University Health Network, Toronto, ON, Canada; Department of Molecular Genetics, University of Toronto, Toronto, ON, Canada; Toronto General Hospital Research Institute, Toronto, ON Canada; The Hull Family Cardiac Fibrillation Management Laboratory, University Health Network Toronto, ON Canada; Department of Laboratory Medicine and Pathobiology, University of Toronto, Toronto, ON, Canada; Institute of Biomedical Engineering, University of Toronto, Toronto, ON, Canada; Division of Cardiac Surgery, Department of Surgery, University of Toronto, Toronto, ON, Canada; Center for Commercialization of Regenerative Medicine (CCRM), Toronto, ON, Canada; King Abdullah University of Science and Technology, Thuwal, Saudi Arabia; Ajmera Transplant Centre, Toronto General Hospital Research Institute, University Health Network, Toronto, ON, Canada

## Abstract

The atrioventricular node (AVN) ensures synchronized heart contractions by establishing the electrical connection between the atria and ventricles. Dysfunction of the pacemaker cells of the AVN leads to atrioventricular block (AV block), a life-threatening condition managed with electronic pacemakers (EPMs). EPMs have drawbacks that could be overcome by a human pluripotent stem cell (hPSC)-derived biological conduction bridge (BioCB). Recent studies demonstrated the differentiation of AVN-like cells from hPSCs, but their conduction properties upon engraftment in vivo remain unexplored. Here we report the generation of AVN-like pacemaker cells (AVNLPCs) from hPSCs using WNT and BMP signaling modulation. These AVNLPCs transcriptionally resemble fetal AVN pacemaker cells, exhibit pacemaker action potentials, and display unique AVN-like conduction properties. Notably, when transplanted into the guinea pig heart, AVNLPCs replicate the functional properties of the AVN. Our study highlights the potential of an AVNLPC-based BioCB as novel cell therapy to improve treatment for AV block patients.

## Introduction

The atrioventricular node (AVN) plays a pivotal role as the electrical bridge between the upper heart chambers (atria) and the bottom heart chambers (ventricles), ensuring synchronized cardiac contractions and efficient blood flow^1–3^. The AVN propagates electrical signals slower than the surrounding myocardium, establishing a conduction delay that allows the ventricles to fill with blood before contracting^4–6^. Additionally, the AVN has the ability to prevent the conduction of excessively fast atrial rates - such as those seen during atrial fibrillation – preventing them from reaching the ventricles and triggering life-threatening ventricular arrhythmias^7–12^. While the sinoatrial node (SAN) serves as the heart’s primary pacemaker^13^, in cases of SAN failure, the AVN can act as a backup pacemaker, initiating ventricular contraction to maintain circulation^14^. Dysfunction of the AVN disrupts impulse transmission, leading to partial or complete AV block (heart block), a condition associated with adverse symptoms ranging from syncope to cardiac death. AV block can be caused by congenital disease (congenital heart block), aging, or cardiac surgery and affects up to 2% of the general population, with prevalence rising to 10% in individuals over 70 years old^15,16^. Currently, AV block is managed through the implantation of electronic pacemakers (EPMs), with ∼1 million devices implanted annually worldwide^17–19^. While EPMs are lifesaving, they have inherent limitations, including the lack of physiological responsiveness to autonomic regulation and risks associated with implantation, such as pacemaker lead complications and infections^20–22^. Additionally, EPMs, particularly those that do not utilize left bundle branch or His bundle pacing, can cause ventricular dyssynchrony, leading to chamber remodeling, fibrosis and heart failure. What is more, EPMs require surgical battery replacements every 10-15 years, which particularly impacts young patients^23–25^. Given the limitations of EPMs, there is a critical need to develop alternative therapies that address the underlying root cause of AVN dysfunction. A promising approach is the development of a human pluripotent stem cell (hPSC)-derived biological conduction bridge (BioCB), as a cell therapy that can restore physiological conduction and provide a regenerative solution for AV block.

In recent years multiple protocols have been developed for the directed differentiation of hPSC-derived atrial, ventricular and pacemaker cardiomyocytes^26–32^. These protocols rely on the successful recapitulation of developmental processes in vitro. The AVN develops from the atrioventricular canal (AVC), a transient embryonic structure between the atria and ventricles that establishes the conduction delay during early heart development. Studies in animal models have demonstrated that AVC specification is driven by WNT signalling, which induces downstream BMP signalling^2,33–35^. BMP signalling, in turn stimulates the expression of TBX2, MSX2, and TBX3, key transcription factors required for AVN cardiomyocyte fate specification and maintenance^33,35,36^. AVC cardiomyocytes either downregulate TBX3 expression and give rise to atrial and ventricular chamber cardiomyocytes or they maintain TBX3 expression and give rise to AVN pacemaker cells^37,38^. Another defining feature of AVN pacemaker cells is the expression of NKX2-5, which distinguishes them from SAN pacemaker cells^39–42^. AVN pacemaker cells can therefore be identified in vitro based on the expression of NKX2-5 and the pan-pacemaker transcription factor TBX3.

In the past two years, protocols utilizing WNT or BMP signalling to differentiate hPSCs into AVC-like cells have been reported^43–46^. These studies performed initial in vitro characterization and demonstrated that AVC-like cells slowly conduct electrical impulses. However, much remains to be learned about their conduction properties and whether they can mimic the key functions of the native AVN. Furthermore, it is still unclear how these cells will function in vivo, which is essential for the development of a BioCB.

Here we generated a NKX2-5^eGFP/wt^TBX3^tdTomato/wt^ reporter hPSC line, enabling precise identification of Atrioventricular Node-like Pacemaker Cells (AVNLPCs) within differentiation cultures. We demonstrate that activation of WNT signalling via the small molecule CHIR99021 triggers downstream activation of BMP signalling, efficiently inducing AVNLPC fate from cardiac progenitors. Interestingly, direct activation of BMP signalling with BMP2 led to more efficient AVNLPC differentiation across multiple hPSC lines. These AVNLPCs transcriptionally match to fetal AVN pacemaker cells and display characteristic pacemaker action potentials. We generated 3D engineered heart tissues and demonstrate that AVNLPCs display slow, decremental conduction properties akin to the endogenous AVN and can effectively block the transmission of arrhythmic impulses. To assess their functional properties in vivo, we used a guinea pig cryoinjury model. In this model, AVNLPCs maintained AVN-like conduction properties for up to 4 weeks in vivo. Importantly AVNLPCs were highly efficient in preventing the conduction of life-threatening arrhythmias in contrast to hPSC-derived ventricular cardiomyocytes that were unable to block these arrhythmias, highlighting the importance of generating the appropriate target cell population for developing a BioCB.

## Results

### NKX2-5^+^ TBX3^+^ cells represent AVNLPCs

To identify NKX2-5^+^TBX3^+^ AVNLPCs in cardiac differentiation cultures we generated a double knock-in NKX2-5^eGFP/wt^ TBX3^tdTomato/wt^ hPSC reporter line^47^ (***Figures 1A* *and S1A-S1D***). Because we previously showed that mesoderm induction conditions impact differentiation efficiencies^29^, we first aimed to identify cardiac mesoderm that would most effectively generate AVNLPCs. To achieve this, we induced RALDH2^+^ atrial mesoderm with 3 ng/ml BMP4 and 2 ng/ml Activin A (3B/2A), and CD235a^+^ ventricular mesoderm with 8 ng/ml BMP4 and 12 ng/ml Activin A (8B/12A), as previously described^29^ (***Figures 1B-1D***). We also included an intermediate induction condition using 5 ng/ml BMP4 and 4 ng/ml Activin A (5B/4A) that resulted in mesoderm expressing 4.9% ± 3.0 RALDH2 and 37.7% ± 7.9 CD235. Notably, the 5B/4A mesoderm yielded the highest proportion of NKX2-5⁺TBX3⁺ cardiomyocytes out of total cells, reaching 31.2% ± 4.0 (***Figure 1E and 1F***). To confirm that NKX2-5⁺TBX3⁺ cells derived from 5B/4A mesoderm represent AVNLPCs, fluorescence-activated cell sorting (FACS) was used to isolate SIRPA^+^CD90^−^ NKX2-5^+^TBX3^+^ and SIRPA^+^CD90^−^NKX2-5^+^TBX3^−^ cardiomyocytes at day 20 and RT-qPCR was performed^48^. SANLPCs (SIRPA^+^CD90^−^NKX2-5^−^TBX3^+^), VLCMs and ALCMs (SIRPA^+^CD90^−^NKX2-5^+^TBX3^−^) were generated from the NKX2-5^eGFP/wt^ TBX3^tdTomato/wt^ hPSC reporter line using previously reported protocols^29,31^ and isolated by FACS at day 20 as gene expression references (***Figures 1G, 1H, S1E, and S1F***). All isolated cell populations expressed comparable levels of the cardiomyocyte marker *TNNT2* validating their cardiomyocyte phenotype. As anticipated, NKX2-5^+^TBX3^+^ cells, VLCMs and ALCMs expressed higher levels of NKX2-5 than SANLPCs. The pacemaker marker *TBX3* was significantly enriched in NKX2-5⁺TBX3⁺ cells and SANLPCs compared to ALCMs, VLCMs, and NKX2-5⁺TBX3⁻ cardiomyocytes derived from the 5B/4A mesoderm induction. The pacemaker ion-channel gene *HCN4* was expressed in NKX2-5⁺TBX3⁺ cells, although at lower levels than in SANLPCs. Importantly, the AVN markers *TBX2*, *BMP2, MSX2* were highly expressed in the NKX2-5^+^TBX3^+^ cells compared to the reference cell types^49^. The SAN marker *SHOX2,* working cardiomyocyte marker *SCN5A*, ventricular markers *MYL2, GJA1* and atrial markers *NPPA, GJA5* were expressed at low levels in the NKX2-5^+^TBX3^+^ cells compared to SANLPC, VLCM and ALCM respectively. These data suggest that NKX2-5^+^TBX3^+^ cells molecularly represent AVNLPCs. To assess the electrophysiological characteristics of FACS-isolated NKX2-5^+^TBX3^+^ cardiomyocytes, action potentials were analyzed via patch clamp (***Figures 1I, 1J and S1G***). NKX2-5^+^TBX3^+^ cells had a spontaneous beating rate of 75±4 beats per minute (bpm) which is comparable to the rate of the human AVN^50,51^ and significantly faster than the rate of VLCMs. In addition, NKX2-5^+^TBX3^+^ cells displayed typical pacemaker action potentials, with significantly slower maximum upstroke velocities (dV/dtmax) compared to VLCM control cells^31,52^. Consistent with a pacemaker phenotype NKX2-5^+^TBX3^+^ cells displayed higher densities of pacemaker funny current (I_f_) compared to VLCMs (***Figure 1K and 1L***). Taken together, these data demonstrate that NKX2-5^+^TBX3^+^ cardiomyocytes have molecular and functional characteristics of AVN pacemaker cells and will be referred to as AVNLPCs hereafter.

**Figure 1:**
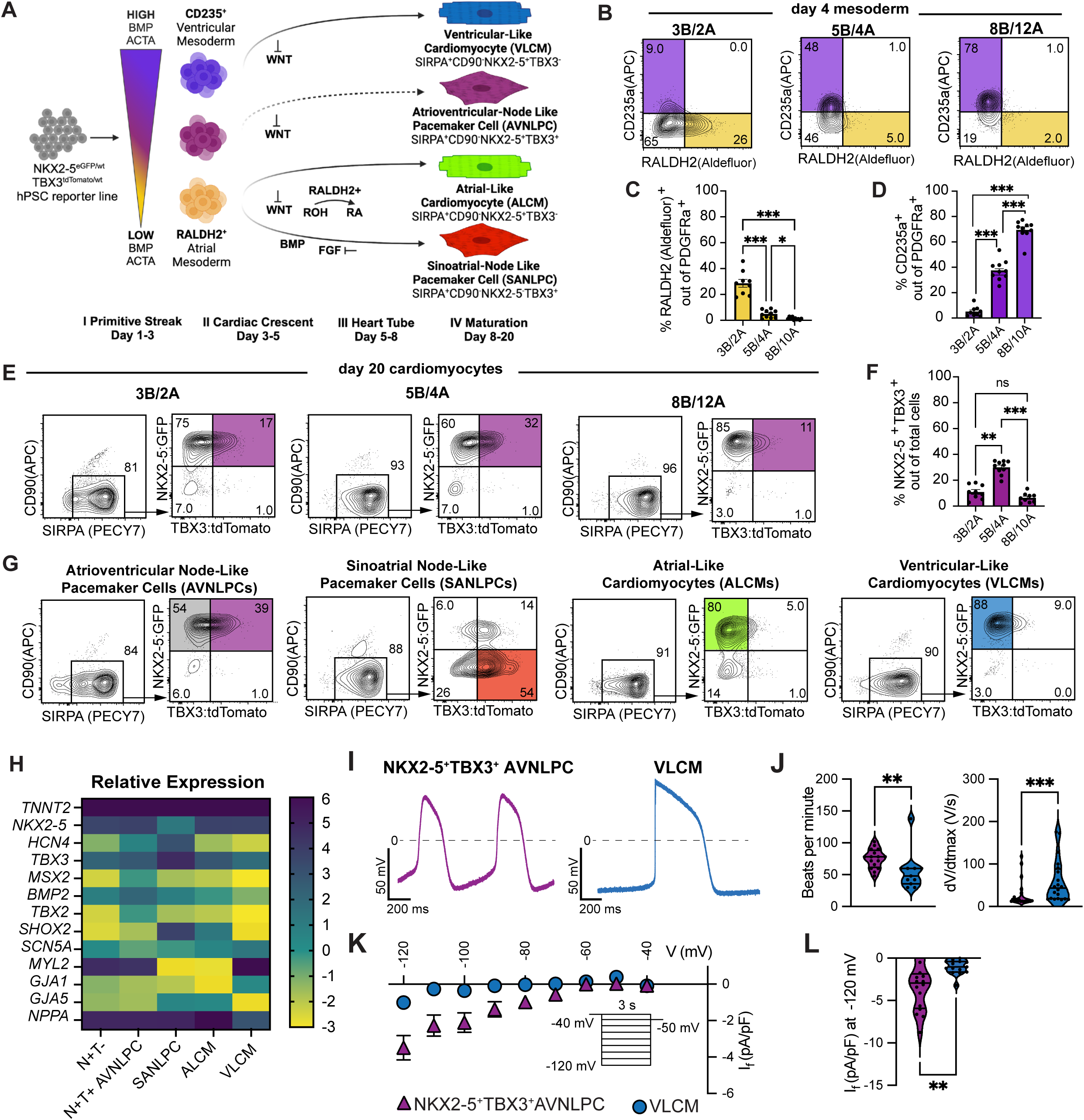
NKX2-5^+^ TBX3^+^ cells represent AVNLPCs. **A.** Summary scheme of developmentally staged cardiomyocyte differentiation protocols showing mesoderm progenitors and respective cardiomyocyte derivatives. High concentrations of Activin A induce a CD235^+^ mesoderm that upon WNT inhibition gives rise to VLCMs. Low concentrations of Activin A induce a RALDH2^+^ mesoderm that upon WNT inhibition and activation of RA signalling gives rise to ALCM. The RALDH2^+^ mesoderm can also give rise to SANLPC following WNT inhibition and activation of RA and BMP signalling and inhibition of FGF signalling. Solid lines: validated lineage relations, dotted lines: proposed AVNLPC lineage relations. **B.** Representative flow cytometric analysis of day 4 mesoderm for the expression of RALDH2 and CD235a after induction with the indicated amounts of BMP4 (B) and Activin A (A) [ng/ml]. **C.** Summary of the proportion of RALDH2^+^ cells in B (n = 10)**. D.** Summary of the proportion of CD235a^+^ cells in B (n=10). **E.** Day 20 representative flow cytometric analysis showing expression of NKX2-5:GFP and TBX3:tdTomato in SIPRA^+^CD90^−^ cardiomyocytes for populations generated with the indicated amounts of BMP4 (B) and Activin A (A) [ng/ml] during mesoderm induction (days1-3). **F.** Summary of the proportion of NKX2-5:GFP^+^TBX3:Tdtomato^+^ cardiomyocytes out of total cells presented in E (n = 10). **G.** Representative flow cytometric plots showing the FACS ating strategy to isolate NKX2-5^+^TBX3^−^ SIRPA^+^CD90^−^ cells (grey) and NKX2-5^+^TBX3^+^SIRPA^+^CD90^−^ AVNLPCs (purple) derived from 5B/4A mesoderm, NKX2-5^−^TBX3^+^SIRPA^+^CD90^−^ SANLPCs (red), NKX2-5^+^TBX3^+^SIRPA^+^CD90^−^ ALCMs (green) and NKX2-5^+^TBX3^+^SIRPA^+^CD90^−^ VLCMs (blue) at day 20 of differentiation from the HES3 double reporter line. **H.** Heat map showing the RT-qPCR gene expression profile of NKX2-5^+^TBX3^+^ AVNLPCs at day 20 of differentiation for the expression of the indicated genes. NKX2-5^+^TBX3^−^SIRPA^+^CD90^−^ sorted cells, SANLPCs, ALCMs, and VLCMs were included as expression reference (n = 7-8). Values represent expression log2 of expression levels relative to the housekeeping gene TBP. **I.** Representative Action potential traces of NKX2-5^+^TBX3^+^ AVNLPCs and VLCMs. **J.** Quantification of indicated action potential parameters. (NKX2-5^+^TBX3^+^ AVNLPCs: n = 16-20, VLCMs: n = 11-20 from three independent differentiations each). **K.** Current-voltage relationship for I_f_ current densities in NKX2-5^+^TBX3^+^ AVNLPCs (n = 16) and VLCMs (n = 15) (inset: voltage protocol). **L.** Violin plot summarizing the maximum I_f_ current densities recorded at -120 mV in NKX2-5^+^TBX3^+^ AVNLPCs (n = 16) and VLCMs (n = 15). Error bars represent SEM. Statistical tests: One-way ANOVA, Bonferroni’s post hoc test (C, D, F); Two-sided t-test (J, L) : *P < 0.05, **P < 0.01, ***P < 0.001. bpm, beats per minute; dv/dt_max_, maximum AP upstroke velocity.

### WNT and BMP signaling pathways guide AVNLPC differentiation

To further increase the differentiation efficiency of the 5B/4A mesoderm into AVNLPCs, we targeted signalling pathways that govern AVN development. AVN cardiomyocytes are specified at the heart tube stage by WNT signaling that induces downstream BMP signaling^33–35,37,53^. To activate WNT signaling the glycogen synthase kinase 3 inhibitor CHIR99021 (CHIR) was administered in 4-day intervals during the Heart Tube to Maturation Stages (days 5-20) (***Figure 2A***). Activation of WNT signalling between days 8-11 and 11-15 significantly increased the proportion of NKX2-5^+^TBX3^+^ AVNLPCs (***Figures 2B and 2C***). Earlier WNT activation (days 5-8) drastically reduced the cardiomyocyte content of the cultures. Using 12 uM CHIR between days 8-11 resulted the highest proportions of AVNLPCs (71.71±8.3% out of total cells, 82.14±7.6% out of SIRPA^+^CD90^+^ cardiomyocytes) and in a 1.5x increase in total cell numbers (CHIR AVNLPCs) (***Figures 2D, 2E, S2A and S2B***). To confirm the activation of WNT target genes and AVN transcription factors following the addition of 12 μM CHIR, RT-qPCR was conducted 3 hours post-treatment. We observed upregulation of *AXIN2* and *LEF1*, along with downstream AVN markers *MSX2*, *TBX2*, and *TBX3* in response to CHIR (***Figure S2C***). Because a relatively high concentration of CHIR was required we tested whether dissociation and reaggregation of cells before CHIR treatment would allow us to lower the concentration (***Figure S2D***). When EBs were dissociated on day 8, 9 μM of CHIR was sufficient to increase AVNLPCs to levels comparable to those achieved with 12 μM CHIR in EBs. To investigate whether the CHIR-induced increase in AVNLPCs is mediated through downstream BMP signaling we inhibited BMP signalling using LDN193189 (LDN) (***Figures S2E and S2F***). When LDN was added together with 12 μM CHIR, the CHIR-induced increase in the proportion of AVNLPCs was significantly reduced to levels comparable to the untreated control. RT-qPCR analysis 3 hours after the addition of the small molecules confirmed that LDN inhibits the CHIR-induced upregulation of BMP target genes *ID1* and *ID2*, thereby blocking the upregulation of AVC genes *MSX2*, *TBX2*, and *TBX*3 (***Figure S2C***).

**Figure 2:**
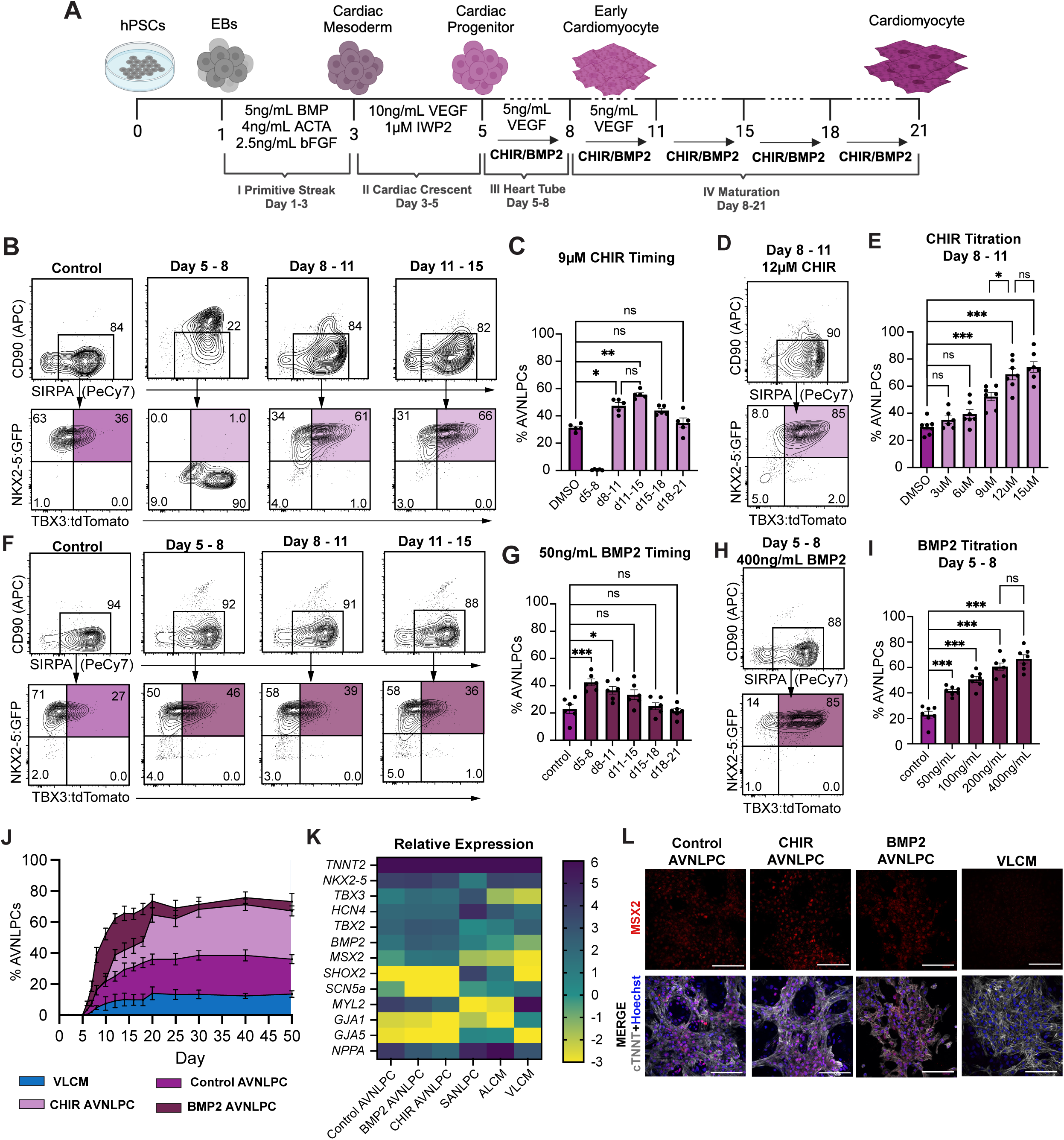
WNT and BMP signaling pathways guide AVNLPC differentiation. **A.** Schematic of the cardiomyocyte differentiation protocol indicating the timepoints when CHIR or BMP2 were added to the cultures. **B.** Representative day 20 flow cytometric analysis of NKX2-5:GFP and TBX3:tdTomato expression in SIPRA^+^CD90^−^ cardiomyocytes after treatment with 9 μM CHIR during the indicated timepoints in comparison to the untreated control. **C.** Bar graph summarizing the proportion of NKX2-5^+^TBX3^+^ cardiomyocytes out of total cells shown in B (n = 5). **D.** Representative day 20 flow cytometric analysis of NKX2-5:GFP and TBX3:tdTomato expression in SIPRA^+^CD90^−^ cardiomyocytes after treatment with 12 μM CHIR at days 8-11 of differentiation. **E.** Proportion of NKX2-5^+^TBX3^+^ cardiomyocytes out of total cells after treatment with the indicated amounts of CHIR at days 8-11 of differentiation. **F.** Representative day 20 flow cytometric analysis of NKX2-5:GFP and TBX3:tdTomato expression in SIPRA^+^CD90^−^ cardiomyocytes after treatment with 50 ng/mL BMP2 during the indicated timepoints in comparison to the untreated control. **G.** Bar graph summarizing the proportion of NKX2-5^+^TBX3^+^ cardiomyocytes out of total cells shown in F (n = 6). **H.** Representative day 20 flow cytometric analysis of NKX2-5:GFP and TBX3:tdTomato expression in SIPRA^+^CD90^−^ cardiomyocytes after treatment with 400 ng/mL BMP2 at days 5-8 of differentiation. **I.** Proportion of NKX2-5^+^TBX3^+^ cardiomyocytes out of total cells after treatment with the indicated amounts BMP2 at days 5-8 during differentiation (n = 7). **J.** Flow cytometric time-course analyses of the proportion of NKX2-5:GFP^+^TBX3^+^ cardiomyocytes out of total cells at the indicated timepoints for untreated Control AVNLPCs (purple), CHIR AVNLPCs (light pink), BMP2 AVNLPCs (maroon) and VLCMs (blue). **K.** Heat map showing the RT-qPCR gene expression profile of Control AVNLPCs, CHIR AVNLPCs and BMP2 AVNLPCs. SANLPCs, ALCMs and VLCMs are included as expression reference. All populations were isolated at day 20 of differentiation. Values represent log2 of expression levels relative to the housekeeping gene TBP (n = 7). **L.** Immunostaining of MSX2 in NKX2-5^+^TBX3^−^SIRPA^+^CD90^−^ VLCMs and NKX2-5^+^TBX3^+^SIRPA^+^CD90^−^ Control/CHIR/BMP2 AVNLPCs isolated from day 20 cultures. Cells were counterstained with cTNT to visualize all cardiomyocytes and DAPI to visualize all cells. Scale bars, 100 μm. Error bars represent SEM. Statistical tests: One-way ANOVA, Bonferroni’s post hoc test (C, E, G, I): *P < 0.05, **P < 0.01, ***P < 0.001. CHIR, CHIR99021

Based on these results, we tested whether activation of BMP signalling could increase the proportion of AVNLPCs. Addition of 400 ng/ml BMP2 during the heart tube stage (days 5-8) resulted in 66.79±8.5% AVNLPCs out of total cells and 75.19±8.9% AVNLPCs out of SIRPA^+^CD90^+^ cardiomyocytes (BMP2 AVNLPCs), comparable to the proportions achieved by WNT activation (***Figures 2F* *-2I and S2G***). In contrast to the WNT activation experiments, total cell numbers were unchanged (***Figure S2H***). RT-qPCR analysis 3 hours after BMP2 addition showed no change in *AXIN2*, *LEF1* or *BMP2* expression but confirmed upregulation of BMP target genes (*ID1, ID2*) and AVN-associated genes (*MSX2, TBX2, TBX3*) compared to untreated control (***Figure S2I***). Dissociation of the cells before treatment did not reduce the required BMP2 concentration (***Figure S2J***). We next tested whether inhibition of BMP signaling could block the development of the baseline proportion of NKX2-5^+^TBX3^+^ AVNLPCs observed in the control cultures. Addition of LDN from days 5-8 resulted in decreased expression of BMP target genes and AVN genes immediately after treatment (3h) and abolished the AVNLPC population at day 20, suggesting that their development is driven by endogenous BMP signalling (***Figures S2I, S2K and S2L***).

NOTCH signaling has been shown to define the boundary between the AVC and the chambers by repressing BMP2 in the atrial and ventricular chambers^54,55^. We therefore reasoned that inhibition of NOTCH signalling might support the development of AVNLPCs. However, addition of the gamma-secretase inhibitor L-685,458 (GSI) at any timepoint (days 3-21) during the differentiation did not increase the proportion of AVNLPCs (***Figure S3A***). Furthermore, other protocols^43,45,46^ that reported differentiation of AVC cells from hPSCs included activation of retinoic acid signalling. Addition of all-trans retinoic acid (RA) at day 3 significantly reduced the proportion baseline AVNLPCs generated in our control condition (***Figure S3B***). This effect was less pronounced when RA (day 3) was added in combination with the CHIR (day 8) or BMP2 (day 5) treatment but still resulted in a significant reduction of AVNLPCs starting at 0.5 μM RA. Based on these results, and the fact that only a small proportion of the 5B/4A mesoderm expressed RALDH2 (<5%), a marker of progenitors reliant on RA signalling for specification, we excluded RA activation in our protocol.

To investigate the timing of NKX2-5 and TBX3 upregulation during the AVNLPC differentiation, a time course analysis was conducted (***Figure 2J***). NKX2-5^+^TBX3^+^ cells were first detected on day 6 of differentiation. The proportion of NKX2-5^+^TBX3^+^ cells increased following CHIR or BMP2 treatment, reaching 64.6±13.2% and 73.5±12.7% respectively by day 20. In contrast, untreated control and VLCM differentiations showed an increase in NKX2-5⁺TBX3⁺ cells, but only to 35.9±9.8% and 14.1±8.3%, respectively. Importantly, the proportion of NKX2-5^+^TBX3^+^ remained high in the CHIR AVNLPC and BMP2 AVNLPC samples and low in the control and VLCM samples until day 50, suggesting a stable phenotype.

To confirm the AVNLPC phenotype of the CHIR AVNLPCs and BMP2 AVNLPCs, we isolated SIRPA^+^CD90^−^NKX2-5^+^TBX3^+^ cardiomyocytes by FACS at day 20 and performed RT-qPCR (***Figures 2K, S3C, and S3D***). SIRPA^+^CD90^−^NKX2-5^+^TBX3^+^ AVNLPCs from an untreated control along with SANLPCs, ALCMs and VLCMs were included as expression references. All samples expressed comparable levels of *TNNT2* and *NKX2-5,* except SANLPCs that had low NKX2-5 expression as expected. CHIR AVNLPCs showed increased expression of the pacemaker marker *TBX3* whereas BMP2 AVNLPCs remained comparable to Control AVNLPCs. Both CHIR and BMP2 AVNLPCs displayed increased expression of *MSX2* in comparison to the untreated Control but had comparable levels of *BMP2 and TBX2. HCN4* was expressed by NKX2-5^+^TBX3^+^ cells in all conditions but at lower levels than in SANLPCs. Expression of the SAN marker *SHOX2* remained low. Expression of ventricular (*MYL*2), atrial (*NPPA, GJA5*) and working cardiomyocyte (*SCN5A* and *GJA1*) genes remained lower in CHIR AVNLPCs and BMP2 AVNLPCs than in VLCMs and ALCMs. Immunostaining confirmed these expression patterns showing that CHIR AVNLPCs and BMP2 AVNLPCs were MSX2^high^ in comparison to untreated control and VLCMs (***Figure 2L***). Finally, we reasoned that combined treatment of BMP2 and CHIR might further improve the AVNLPC differentiation. However, the proportion of AVNLPCs and the expression of AVN marker genes was comparable between CHIR only, BMP2 only and BMP2+CHIR treated samples (***Figures S3E-S3G***). Collectively, these findings demonstrate that CHIR or BMP2 treatment is sufficient to differentiate hPSCs into AVNLPCs.

### Generation of AVNLPCs from other hPSC lines

To test the applicability of the protocol to other hPSC lines, we differentiated AVNLPCs from the HES2 and ESI-017 ASAP1 lines^56,57^. Since endogenous signaling varies between cell lines, BMP4 and ACTA were titrated to determine which induction condition results in a mesoderm that efficiently gives rise to AVNLPCs. We aimed at generating a mesoderm that is ∼40% PDGFRa^+^CD235a^+^ at day 4, as this mesoderm generated the highest proportion of NKX2-5^+^TBX3^+^ cells in the NKX2.5^eGFP/wt^TBX3^tdTomato/wt^ reporter line. For the HES2 and ESI-017 lines induction with 5B/2A resulted in mesoderm expressing 31.85±6.2% and 37.07±12.5% CD3235a respectively (***Figures 3A and 3B***). Induction with higher concentrations of cytokines (8B/12A, 8B/10B respectively) resulted in ventricular-like mesoderm expressing high levels of CD235a. Both cell lines efficiently generated cardiomyocytes with lower proportions of MLC2v^+^ cells obtained from the 5B/2A inductions (***Figures 3C and 3D***). Next, the differentiations were optimized to efficiently produce cTNNT^+^ CHIR AVNLPCs and BMP2 AVNLPCs (***Figures 3E-3H***). In contrast to the HES3 reporter hPSC line, the proportion of cardiomyocytes significantly improved when the CHIR and BMP2 treatment windows were delayed by 3 days (day 11-15 and day 8-11 respectively). In addition, significantly higher cardiomyocyte proportions were observed in BMP2 AVNLPC cultures compared to CHIR AVNLPC cultures. Therefore, magnetic activated cell sorting (MACs) using the PSC-derived Cardiomyocyte Isolation Kit was performed prior to RT-qPCR expression analysis to equalize the cardiomyocyte content between samples. CHIR AVNLPCs and BMP2 AVNLPCs from the HES2 and ESI-017 cell line express AVN markers (*TBX3, HCN4, MSX2, BMP2, TBX2*) at comparable levels to the HES3-derived AVNLPCs and at higher levels than VLCMs (***Figure 3I***). Ventricular and Atrial genes were expressed at low levels like in the HES3-derived AVNLPCs. To further quantify the differentiation efficiency in the BMP2 AVNLPC cultures, we performed immunofluorescent staining for MSX2 which showed 59.71±14.59% and 61.06±14.72% MSX2^+^ AVNLPCs for the HES2 and the ESI017 cell line respectively (***Figures 3J and 3K***). Taken together, these data show that our protocols allow for the efficient generation of AVNLPCs from multiple hPSC lines. Due to the negative impact of the CHIR treatment on the overall cardiomyocyte content we decided to focus all remaining experiments on AVNLPCs generated with the BMP2 protocol.

**Figure 3:**
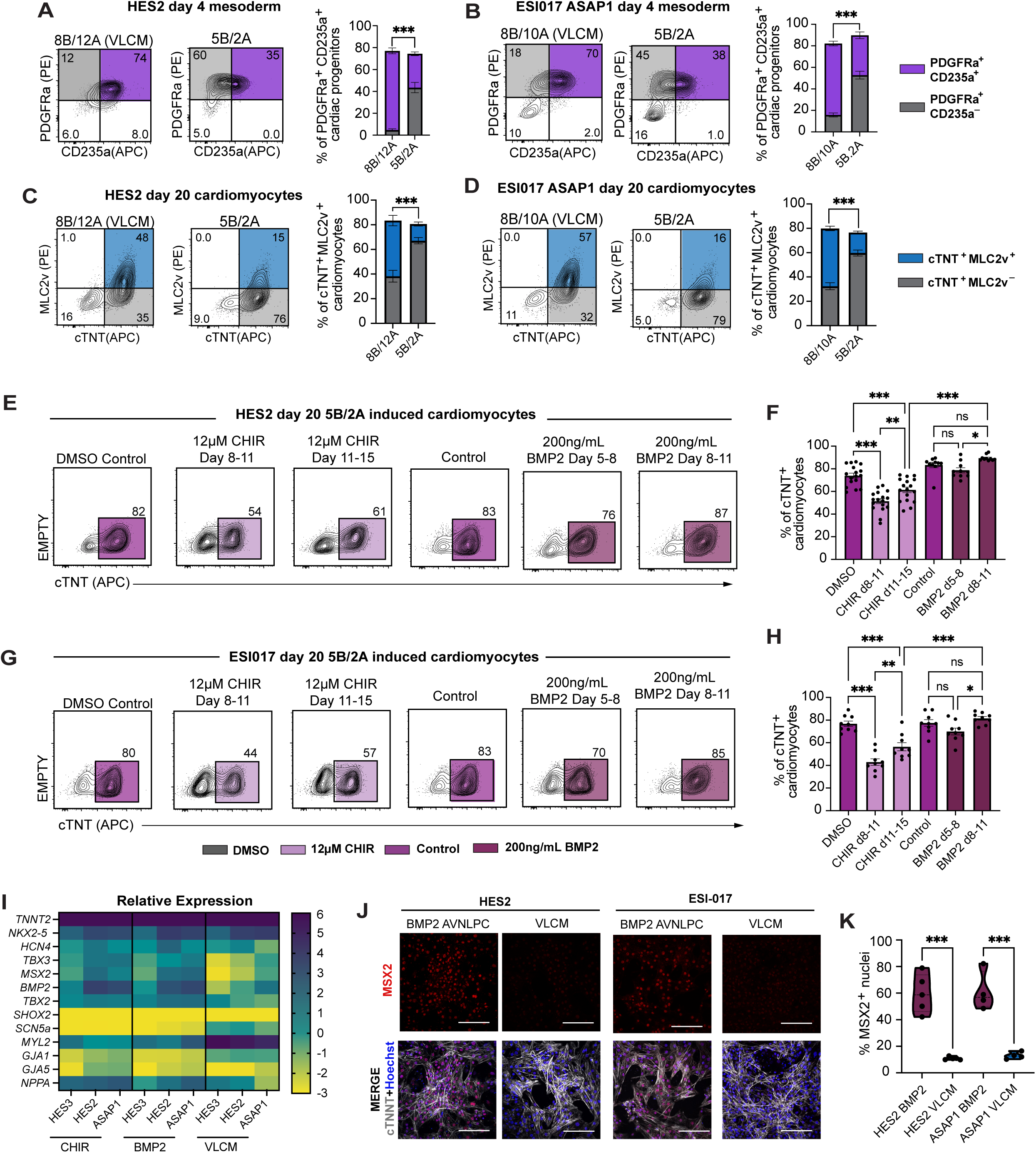
Generation of AVNLPCs from other hPSC lines. **A.** Representative flow cytometric analysis and corresponding summary of CD235a and PDGFRα expression in day 4 mesoderm following induction with the specified concentrations of BMP4 (B) and Activin A (A) [ng/ml] in the HES2 cell line (n = 14). **B.** Representative flow cytometric analysis and corresponding summary of CD235a and PDGFRα expression in day 4 mesoderm following induction with the specified concentrations of BMP4 (B) and Activin A (A) [ng/ml] in the ESI017 ASAP1 cell line (n = 15-16). **C.** Representative flow cytometric analysis and corresponding summary of cTNT and MLC2V expression in day 20 populations generated with the indicated amounts of BMP4 (B) and Activin A (A) [ng/ml] during mesoderm induction (days 1-3) in the HES2 cell line (n = 14). **D.** Representative flow cytometric analysis and corresponding summary of cTNT and MLC2V expression in day 20 populations generated with the indicated amounts of BMP4 (B) and Activin A (A) [ng/ml] during mesoderm induction (days 1-3) in the ESI017 ASAP1 cell line (n = 15-16). **E.** Representative flow cytometric analysis for the expression of cTNT in day 20 cultures treated with either 12 μM of CHIR or 200 ng/mL of BMP2 at the indicated timepoints in the HES2 cell line. **F.** Bar graph summarizing the proportion of cTNT^+^ cells shown in F (n = 14). **G.** Representative flow cytometric analysis for the expression of cTNT in day 20 cultures treated with either 12 μM of CHIR or 200 ng/mL of BMP2 at the indicated timepoints in the ESI-017 ASAP1 cell line. **H.** Bar graph summarizing the the proportion of cTNT^+^ cells shown in G (n = 9). **I.** Heatmap showing differences in RT-qPCR gene expression levels across HES3, HES2 and ASAP1 AVNLPCs treated with either 12 μM of CHIR (CHIR AVNLPCs) or 200 ng/mL of BMP2 (BMP2 AVNLPCs). VLCMs differentiated from each cell line were included as an expression reference. All populations were analyzed at day 20 of differentiation. Values represent log2 of expression levels relative to the housekeeping gene TBP. **J.** Immunofluorescent staining of MSX2 in BMP2 AVNLPCs and VLCMs from the HES2 and ESI-017 ASAP1 cell lines. Cells were counterstained with cTNT to visualize all cardiomyocytes and DAPI to visualize all cells. Scale bars, 100 μm. **K.** Quantification of the percentage of MSX2^+^ cells determined by immunofluorescence staining in BMP2 AVNLPCs and VLCMs from the HES2 and ESI-017 ASAP1 cell lines (n = 4-5 images per monolayer, taken from 3 independent differentiations). Error bars represent SEM. Statistical tests: One-way ANOVA, Bonferroni’s post hoc test (F, H); unpaired two-sided t-test (A-D, K): *P < 0.05, **P < 0.01, ***P < 0.001.

### Establishing a single-cell transcriptome expression reference of the fetal AVN

We performed single-nucleus RNA sequencing (snRNA-seq) of a 19-week gestation fetal heart to establish a fetal AVN gene expression reference for comparison with our in vitro AVNLPCs (***Figure 4A***). To increase the number of AVN pacemaker cells that are included in this analysis we specifically dissected the tissue of the Atrioventricular junction. Unsupervised clustering identified 9 cell clusters, encompassing expected cell types, including non-cardiomyocytes (fibroblasts, smooth muscle, endothelial and endocardial cells) and cardiomyocytes (***Figure 4B, Table S1 and S2***). As discussed, the AVN is a heterogeneous structure composed of transitional cells, with the Core AVN at its center. To resolve this complexity, we focused our analysis on the cardiomyocyte population within the dataset. Subclustering the *TNNT2* expressing cardiomyocytes yielded 20 additional clusters (***Figures S4A and S4B***). Clusters were merged based on similarity in marker gene expression, resulting in the identification of seven major cell types: Core AVN, Atrial Transition Zone (ATZ), Lower Nodal Bundle (LNB), His Bundle, ventricular cardiomyocytes, atrial cardiomyocytes, and proliferating cardiomyocytes (***Figures 4C, S4C, S4D, and Table S3***). Core AVN cells were identified by high expression of AVN-specific genes (*RSPO3, MSX2, BMP2, TBX2*) and pacemaker markers (*TBX3, HCN4, GJC1*) (***Figure 4D***). The Core AVN cells exhibited low expression of fast-conduction and working cardiomyocyte genes (*GJA5, GJA1, SCN5A*) and were negative for SAN pacemaker markers *SHOX2, CD34* and *MYH11*^58^. Surrounding the Core AVN are transitional cells with gradually shifting gene expression. Accordingly, ATZ cells were defined by co-expression of AVN and pacemaker genes (*RSPO3, BMP2, TBX3, HCN4, GJC1*), alongside atrial markers (*NPPA, NR2F2, GJA5*). Similarly, the LNB cells expressed pacemaker genes (*TBX3, GJC1*) alongside ventricular markers (*MYL2, IRX4*) but low levels of fast-conduction and working cardiomyocyte genes (*GJA5, GJA1, SCN5A*). A cluster expressing AVN markers comparable to the Core AVN, along with fast conducting ion-channels *GJA5 and SCN5A* was annotated as the His Bundle. Two cell clusters representing atrial cells were identified and merged based on the expression of atrial markers *NPPA* and working myocyte markers *GJA5*, *GJA1* and *SCN5A.* Furthermore, nine ventricular clusters were identified and merged based on the expression of *MYL2*, *IRX4*, *GJA1*, and *SCN5A*. Lastly, five clusters of proliferating *MKI67*-expressing cells were identified. Cell type annotation was further validated through gene signature scoring using marker gene sets from a recently published scRNA-seq dataset of the developing human heart by Farah et al. (***Figure 4E***)^59^. This dataset collected earlier stages of fetal heart development (gestation week 9-15) and accordingly identified an AVC cluster. Our AVN and ATZ clusters scored the highest with the Farah et al. AVC cluster gene set confirming our cluster annotation. Similarly, the identity of our Atrial, Ventricular and His bundle clusters were confirmed based on scoring with the differentially expressed genes (DEGs) of the respective subtypes reported by Farah et al. In addition, DEGs from adult heart AVN cells identified by Kanemaru et al.^42^ scored strongly to our fetal AVN cluster, further supporting its identity (***Figure 4F***).

**Figure 4:**
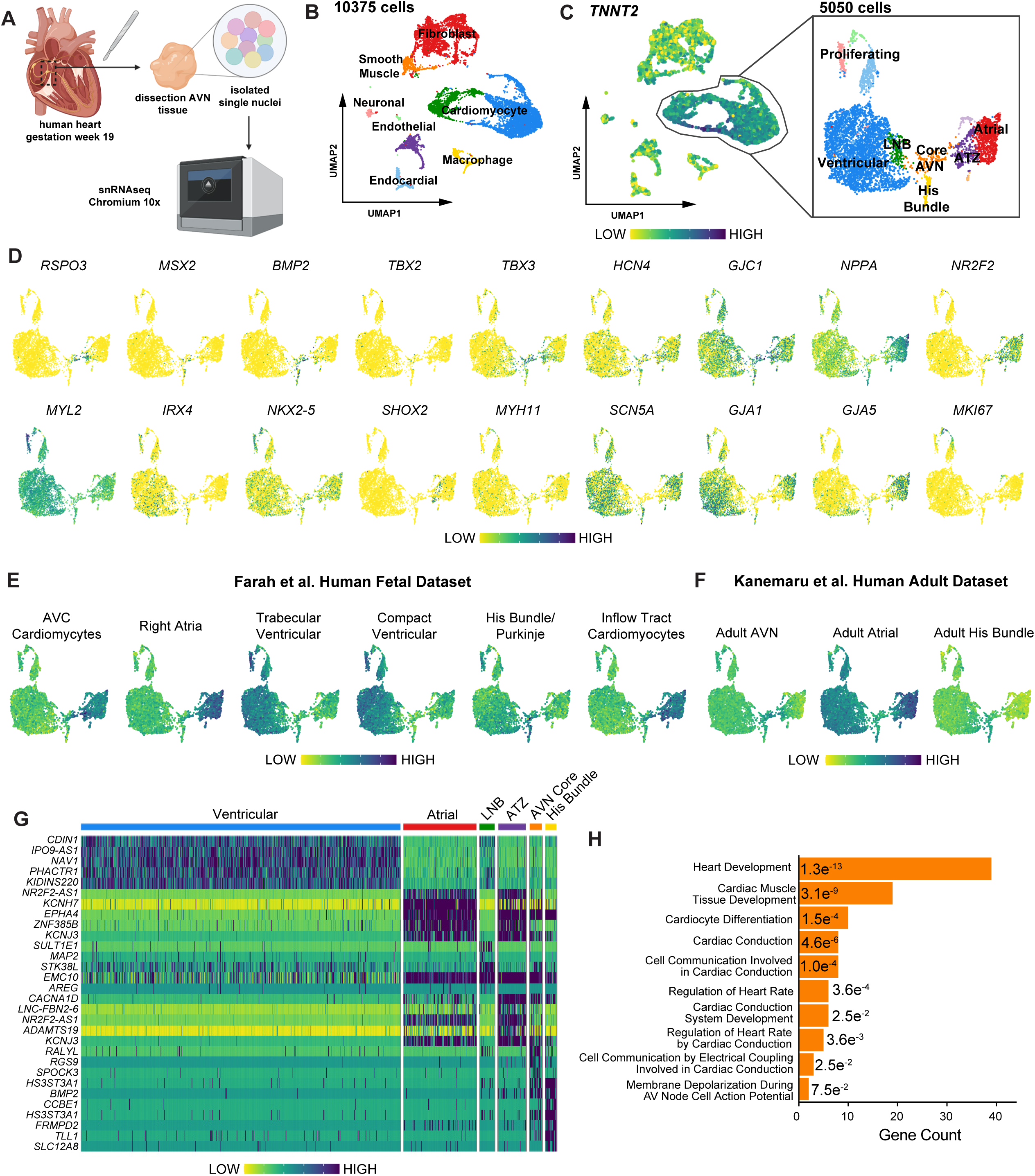
Single-nucleus transcriptomic profiling of the fetal AVN. **A.** Schematic of the dissection and preparation of AVN tissue for single-nucleus RNA sequencing (snRNA-seq) using the 10x Genomics Chromium platform. **B.** Uniform manifold approximation and projection (UMAP) of gestation week 19 fetal heart AVN tissue identifying 9 cell clusters. **C.** UMAP depicting TNNT2^+^ clusters (left). Sub clustering of the *TNNT2*^+^ cardiomyocytes showing 7 subclusters (right). **D.** UMAPs of the subclustered cardiomyocyets depicting the expression of the indicated genes. **E.** UMAPs showing signature score distribution for the DEGs of indicated fetal heart cell types from Farah et al. spatial MERFISH dataset. **F.** UMAPs showing signature score distribution for the DEGs of indicated adult heart cell types from Kanemaru et al. dataset. **G.** Heatmap displaying the top 5 DEGs for each cardiomyocyte subcluster. **H.** GO enrichment analysis of biological processes associated with the positive differentially expessed genes in the Core AVN cluster.

The top DEGs were identified for each cardiomyocyte subtype (***Figure 4G* *and Table S3***). The top 10 DEGs in the Core AVN cluster included several genes also enriched in the AVC cluster from the Farah et al. dataset, such as *RGS9, BMP2, TBX3 and RSPO3* (***Figure S4E***)^59^. Gene ontology (GO) analysis of biological processes further supported the AVN identity of the Core AVN cluster, with enriched genes associated with terms such as cardiac conduction, cell communication involved in cardiac conduction, heart rate regulation, cardiac conduction system development, and membrane depolarization during AV node cell action potential (***Figure 4H***).

### AVNLPCs resemble fetal Core AVN pacemaker cells

Next, we leveraged our fetal AVN dataset to benchmark our hPSC-derived BMP2 AVNLPCs and assess their transcriptional similarity to endogenous AVN cells. Single-cell RNA sequencing (scRNA-seq) was performed on day 20 cells derived from the HES2 hPSC line using the optimized BMP2 AVNLPC differentiation protocol (***Figure 5A***). Unsupervised clustering revealed five distinct cell clusters with the majority of cells representing *TNNT2^+^* cardiomyocytes alongside small clusters of endothelial cells, fibroblasts and epicardial cells (***Figure 5B, Tables S1 and S4***). Spearman correlation analysis showed that the *TNNT2*^+^ cells contained in the BMP2 AVNLPC cultures strongly correlated with the fetal Core AVN (***Figure 5C***). GO analysis of biological processes further supported the AVN identity of the BMP2 AVNLPCs, linking enriched genes in the cardiomyocyte cluster to terms like regulation of heart contraction, cardiac conduction, cardiac pacemaker cell differentiation, AV node cell action potential and atrioventricular node cell differentiation (***Figure 5D***). Thus, single-cell transcriptome analysis confirmed that our BMP2 AVNLPC differentiation protocol generates cardiomyocytes with an AVN phenotype, resembling the cells of the fetal Core AVN.

**Figure 5:**
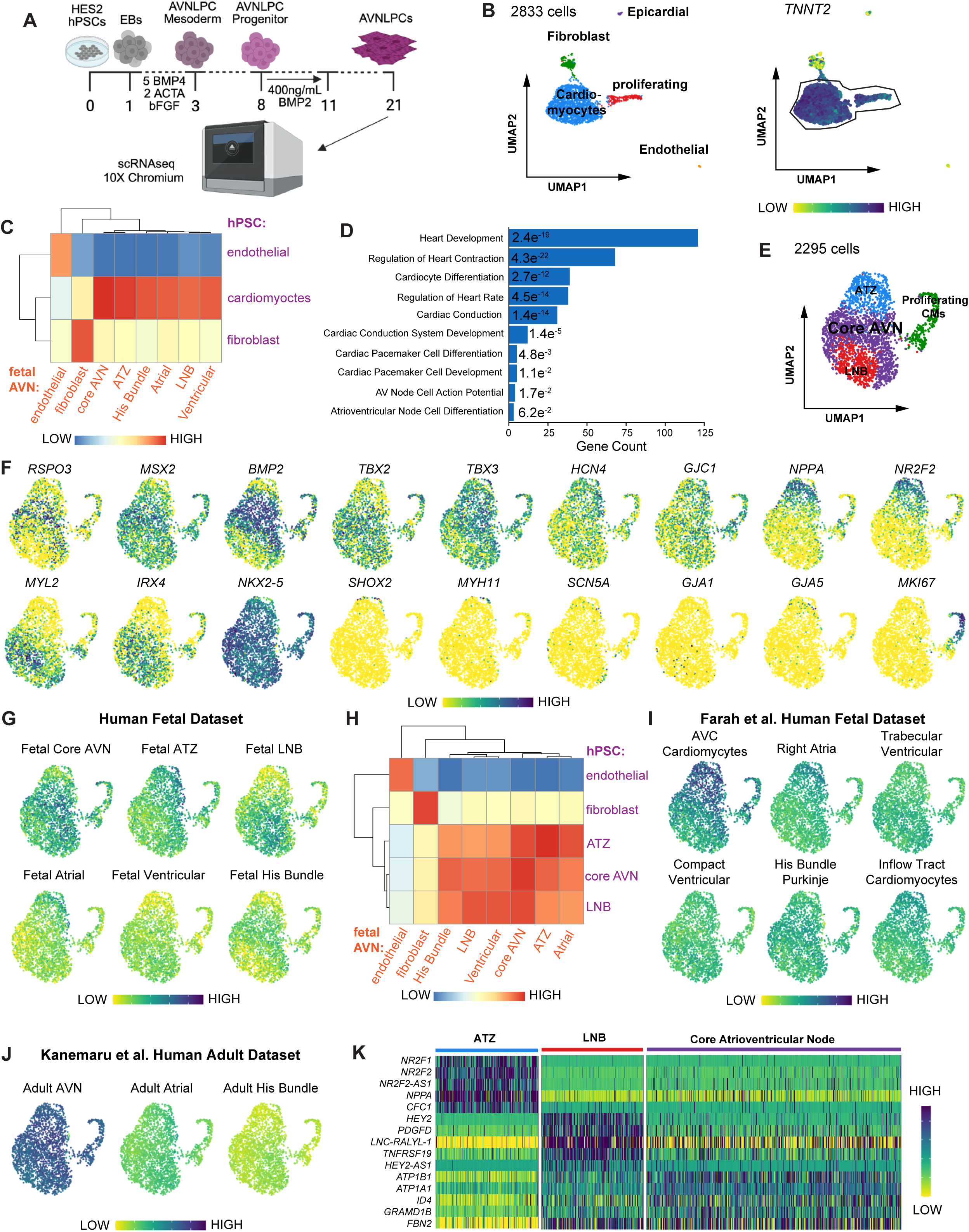
Single-cell RNA sequencing of hPSC-derived AVNLPCs reveals transcriptomic similarities to fetal AVN cardiomyocytes. **A.** Schematic of the BMP2 AVNLPCs differentiation protocol in the HES2 cell line and subsequent processing on the 10x Genomics Chromium platform for scRNA-seq. **B.** UMAP of day 25 hPSC-derived AVNLPCs showing 5 cell clusters (left). UMAP depicting TNNT2^+^ clusters (right). **C.** Spearman correlation between selected cell clusters from the fetal AVN and hPSC-derived AVNLPC dataset. **D.** GO enrichment analysis of biological processes associated with the positive differentially expessed genes in the hPSC-derived AVNLPC cardiomyocyte cluster. **E.** UMAP of subclustered *TNNT2***^+^** cardiomyocytes showing 4 subclusters. **F.** UMAPs of the subclustered cardiomyocytes depicting the expression of the indicated genes. **G.** UMAPs showing signature score distribution for the DEGs of the indicated fetal AVN cell types. **H.** Spearman correlation between selected cell clusters from the fetal AVN and hPSC-derived AVNLPC datasets, p < 0.05 for all correlations. **I.** UMAPs showing signature score distribution for the DEGs of indicated fetal heart cell types from the Farah et al. spatial MERFISH dataset. **J.** UMAPs showing signature score distribution for the DEGs of indicated adult heart cell types from the Kanemaru et al. dataset. **K.** Heatmap showing the top 5 differentially expressed genes of the AVNLPC cardiomyocyte subtypes.

We next wanted to see whether the in vitro cultures also contain the different AVN cardiomyocyte subtypes that we identified in the fetal heart. To this end we subclustered the *TNNT2*^+^ cells and identified seven distinct clusters (***Figures S5A and S5B***). Clusters were merged based on similarity in marker gene expression profiles, resulting in the identification of 4 major cell types: AVN core, LNB, ATZ, and proliferating cardiomyocytes (***Figures 5E, S5C, S5D and Table S5***). Three clusters of Core AVN cells were identified and merged based on exhibiting the highest expression of key AVN markers, including *RSPO3*, *MSX2*, *BMP2*, and *TBX2.* (***Figure 5F***). The ATZ cluster expressed lower levels of these AVN genes together with elevated levels of atrial markers *NPPA and NR2F2.* The LNB cluster displayed reduced expression of AVN and pacemaker markers, along with elevated expression of ventricular markers *MYL2* and *IRX4*. All *TNNT2*^+^ cardiomyocyte clusters were *NKX2-5*^+^, *SHOX2^−^, and MYH11*^−^, confirming their AVN rather than SAN pacemaker identity. None of the clusters exhibited high expression of the working cardiomyocyte markers *SCN5A, GJA1 and GJA5* most likely due to the AVN-like phenotype of all the cardiomyocytes subtypes contained in the BMP2 AVNLPC cultures. Additionally, two clusters of proliferating cardiomyocytes were identified, marked by *MKI67* expression.

To further validate the identity of the Core AVN, ATZ, and LNB clusters we performed gene signature scoring of our hPSC dataset with the DEG lists from our fetal tissue dataset (***Figure 5G***). The fetal core AVN DEGs scored the highest in the three cell clusters making up the Core AVN of the hPSC dataset. Similarly, the ATZ and LNB cell clusters from the hPSC dataset were found to score most strongly with the DEGs from the fetal ATZ and fetal LNB, respectively. This analysis also confirmed that the BMP2 AVNLPCs in the hPSC dataset do not strongly correlate to fetal Ventricular, Atrial or His Bundle cells. Spearman correlation analysis further demonstrated that the hPSC-derived Core AVN cells closely resemble fetal Core AVN cells (***Figure 5H***).

For additional validation of the AVN phenotype we also compared the hPSC dataset to published fetal heart (Farah et al.)^59^ and adult heart (Kanemaru et al.)^42^ datasets (***Figures 5I and 5J***). Gene signature scoring of the fetal heart dataset showed highest correlation of our BMP2 AVNLPCs with fetal AVC cardiomyocytes while the other cardiomyocyte subtypes (Right Atrial, Ventricular, His Bundle and Purkinje, Inflow Tract) contained in this dataset resulted in lower scores. Similarly, adult heart AVN cell DEGs scored highest in our BMP2 AVNLPCs in contrast to low scores for Adult Atrial and His Bundle DEGs. Taken together, this single-cell transcriptome analysis further confirms the AVN identity of our BMP2 AVNLPCs and demonstrates that the cultures contain subtypes of AVN pacemaker cells including Core AVN and Transition Zone cells similar to the fetal AVN. Finally, DEG analysis of the hPSC-derived Core AVN, ATZ, and LNB cells highlighted the gene expression patterns in these cardiomyocyte subtypes (***Figure 5K* *and Table S5***). The top 10 DEGs in the Core AVN cluster included several genes also enriched in our fetal Core AVN cluster, such as *ATP1B1, ID4, GRAMD1B, FBN2 and BMP2 (**Figure S5E**)*.

### Engineered tissues containing AVNLPCs mimic conduction properties of the AVN

To assess the electrophysiological characteristics of AVNLPCs, we took advantage of the genetically encoded ASAP1 voltage sensor of the ESI-017-ASAP1 cell line^56^ allowing for analysis of electrical activity via optical mapping^60–62^. We isolated ESI-017 ASAP1-derived AVNLPCs and VLCMs using the PSC-derived Cardiomyocyte Isolation MACs Kit to achieve a standardized cardiomyocyte content of >80%. Purified AVNLPCs or VLCMs were seeded together with cardiac fibroblasts in a 3:1 ratio into molds to make 3 mm long engineered AVN tissues (eAVNT) or engineered ventricular tissues (eVT) respectively (***Figures 6A and 6B***)^63^. We first recorded spontaneous optical action potentials (***Figures 6C and 6D***). As expected for pacemaker cells, the eAVNTs displayed a steady increase in membrane potential during phase 4 of the action potential, known as diastolic depolarization^52,64^. In comparison, eVTs showed significantly lower rates of diastolic depolarization. eAVNTs further displayed significantly slower maximum action potential upstroke velocities compared to eVTs. The action potential durations at 30% (APD30) and 90% (APD90) of repolarization was not significantly different between eAVNTs and eVTs (***Figure S6A***). Since the AVN acts as the conduction bridge between the atria and the ventricles and establishes a conduction delay, we next analyzed the conduction properties of the engineered tissues while pacing them from the bottom at 2.5 Hz (***Figures 6E and 6F***). The eAVNTs had slow conduction velocities (3.8±1.5 cm/s) comparable to the human AVN^1^. In contrast, eVTs had significantly faster conduction velocities (12.8±2.8 cm/s). An important function of the AVN is to block fast atrial rhythms, such as those seen in atrial fibrillation, from reaching the ventricles and causing ventricular tachycardia, which can result in sudden cardiac death. To assess this function, we simulated fast atrial rhythms by increasing the pacing frequency (***Figures 6G-6I***). eAVNTs showed a 1:1 capture ratio until 3.3±0.2 Hz, a physiological equivalent of 198 beats per minute (bpm). At higher frequencies eAVNTs transitioned into a 2:1 conduction block and eventually a 4:1 conduction block. In contrast, eVTs showed a 1:1 capture ratio up to 4.8±0.3 Hz, equivalent to 288 bpm. These data suggest that eAVNTs can more effectively block fast atrial rhythms than eVTs. We next assessed the underlying mechanisms of this conduction property. 100% of eAVNTs exhibit the AVN-specific Wenckebach Phenomenon, which is defined as a progressive increase of impulse conduction time (S1-R1) at a fixed stimulation frequency (S1-S1), leading up to a non-conducted impulse, or blocked beat^8,9^ (***Figures 6J, 6K and S6B***). In contrast the majority (73%) of eVTs did not show a progressive increase in impulse conduction before blocking a beat when paced at a fixed frequency. We next used a S1-S2 stimulation protocol to assess whether eAVNTs display decremental conduction like the endogenous AVN^11,12^ (***Figures 6L, S6C and S6D***). In this protocol we applied 8 stimuli at a stable frequency (S1) of 2.5Hz followed by an extra stimulation (S2) with an increasing frequency from 3Hz to 10Hz^65–67^. In eAVNT tissues the impulse conduction time from S2 stimulation to the activation of the action potential (R2) progressively increased with increasing S2 frequency, characteristic of decremental conduction. This impulse conduction time, also referred to as latency, increased until conduction of the extra beat was blocked at 3.2±0.1 Hz. A slow frequency-dependent increase in latency was also observed in the eVTs, with S2 stimulations being blocked 4.1±0.4 Hz. The slope of the latency increase was significantly steeper in eAVNTs compared to eVTs indicating stronger decremental conduction in eAVNTs. Immunofluorescence staining of the tissues showed that they maintained their molecular identify with eAVNTs staining NKX2-5^+^ MSX2^high^, whereas eVTs stained NKX2-5^+^ MSX2^low^ (***Figures 6M, and S6E***). We repeated this conduction property analysis in the HES3 NKX2.5^eGFP/wt^TBX3^tdTomato/wt^ reporter line which confirmed the results described above (***Figures S6F-S6H***). Taken together, these data demonstrate that engineered 3D tissues containing AVNLPCs display action potential characteristics and conduction properties like the endogenous AVN.

**Figure 6:**
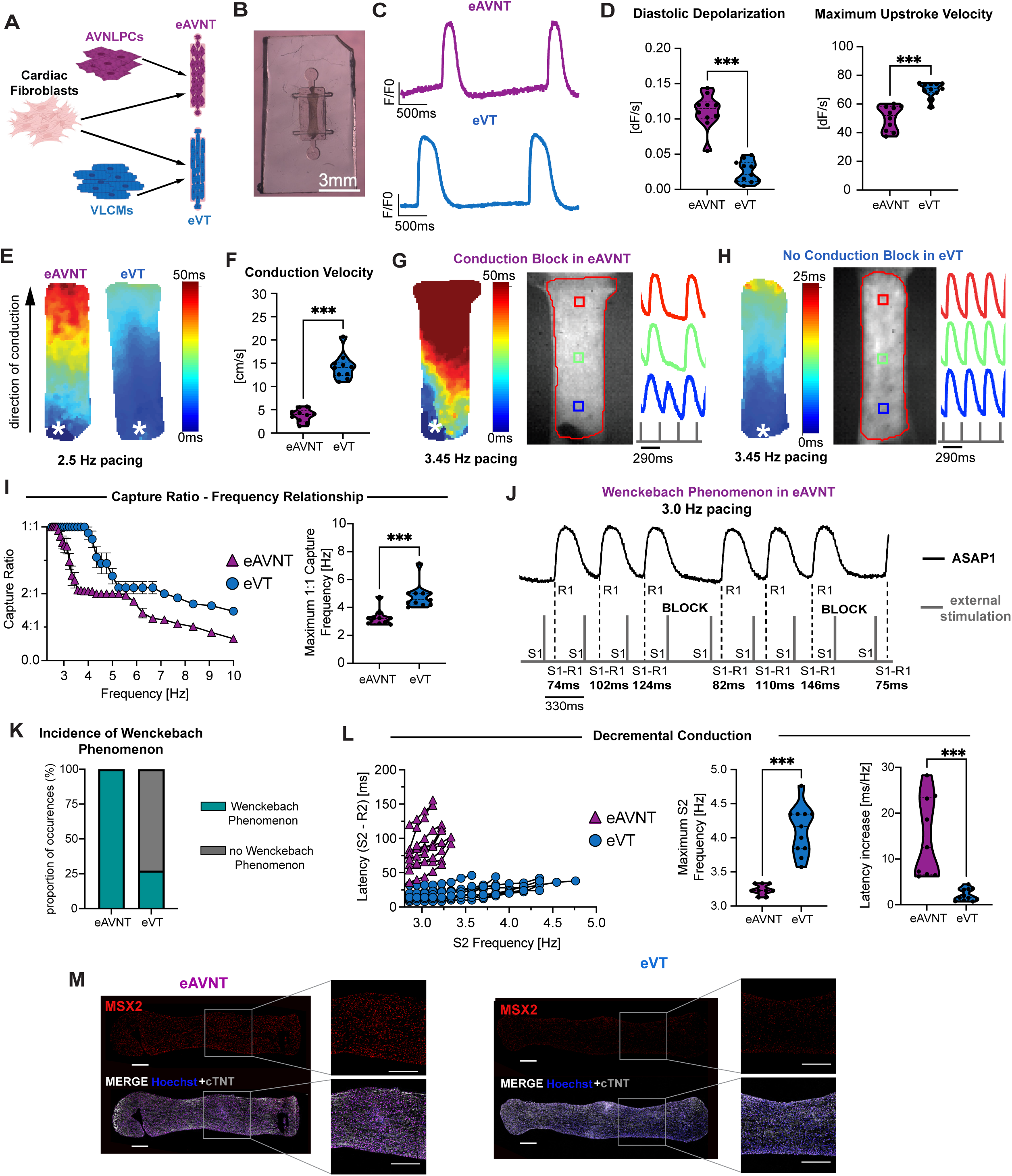
Engineered tissues containing AVNLPCs mimic conduction properties of the AVN. **A.** Schematic illustrating the process of combining cardiac fibroblasts with AVNLPCs or VLCMs to generate engineered heart tissues referred to as eAVNTs and eVTs respectively. **B.** Representative bright field image of an eAVNT. **C.** Representative optical action potential traces recorded in an eAVNT and eVT. **D.** Quantification of indicated optical action potential parameters (eAVNTs: n= 2-3, eVTs: n=2-3, from four independent differentiations each). **E.** Representative voltage activation map of an eAVNT and eVT. White asterisk indicates location of pacing electrode (bottom of engineered tissues). Black arrow indicates the direction of conduction. **F.** Quantification of conduction velocities recorded in eAVNTs and eVTs. (eAVNTs: n = 2-3, eVTs n = 2-3 from four independent differentiations each). **G.** Representative voltage activation map of an eAVNT showing 2:1 conduction block when paced at 3.45 Hz (left). Corresponding brightfield image and optical action potentials near the pacing source (blue), in the middle (green), and at the distal end (red) of the tissue (right). White asterisk indicates location of pacing electrode. **H.** Representative voltage activation map of an eVT not showing a conduction block when paced at 3.45 Hz (left). Corresponding brightfield image and optical action potentials near the pacing source (blue), in the middle (green), and at the distal end (red) of the tissue (right). White asterisk indicates location of pacing electrode. **I.** Graph summarizing the capture ratios of eAVNTs and eVTs at increasing pacing frequencies (left). Quantification of the frequency at which 1:1 capture is lost (right). (eAVNTs: n= 2-3, eVT n=2-3, from four independent differentiations each). **J.** Representative optical action potential trace of an eAVNT displaying the Wenckebach Phenomenon leading up to a blocked beat when paced at 3Hz. S1 (grey) denotes the applied stimulus via the pacing electrode, and R1 represents the tissue’s action potential response to S1. S1-R1 quantifies the impulse conduction time. **K.** Quantification of the proportion of eAVNTs and eVTs displaying the Wenckebach Phenomenon. **L.** Assessment for decremental conduction properties in eAVNTs and eVTs. Graph displaying the response of the impulse conducting time (Latency, S2-R2) to increasing frequency of the extra stimulus (S1-S2) (left). Quantification of the frequency of the extra stimulus (S2) at wich capture is lost (middle). Quantification of the slope of the latency increase (right). (eAVNTs: n = 2-3, eVTs n = 2-3 from four independent differentiations each). **M.** Immunofluorescent staining of MSX2 in eAVNTs and eVTs counterstained with cTNT to identify cardiomyocytes and Hoechst to visualize all cells. Scale bars, 250 μm. Error bars represent SEM. Statistical tests: unpaired two-sided t-test (D, E, G, J, M): **P < 0.01, ***P < 0.001. S1-S2, cycle length of extra stimulus; S2-R2, impulse conduction time from S2 stimulus to action potential response in tissue (R2) = Latency.

### AVNLPCs exhibit AVN conduction properties in vivo

To test whether AVNLPCs maintain their AVN-like conduction properties in vivo we adapted a previously reported guinea pig cryoinjury model^56,68^ (***Figure 7A***). First, we used cryoinjury to induce formation of scar tissue and establish an artificial conduction block in the left ventricular free wall. After ten days, 80-100 million ESI-017-ASAP1-dervied AVNLPCs or VLCMs were injected into the border zone of the healthy myocardium spanning into the transmural scar. We reasoned that this would allow us to assess electrical conduction from the healthy myocardium into the electrically insulated scar tissue via the cell grafts. Assessment of graft conduction within the insulated scar avoids the potential impact of underlying fast conducting guinea pig myocardium, which could otherwise lead to inaccurate conduction measurements within the graft. After two to four weeks post-transplantation the hearts were harvested and mounted to a Langendorff perfusion system to allow for ex vivo electrocardiogram (ECG) recording and optical mapping. The genetically encoded ASAP1 voltage sensor was used to analyze the electrical activity of the hPSC-derived grafts (***Figures 7B, 7C and S7A***). To assess the electrical activity of the host, hearts were perfused with the voltage-sensitive dye RH237^68,69^. To control the heart rate, we perfused the hearts with the muscarinic agonist methacholine, which supresses both SAN and AVN automaticity, effectively silencing the guinea pig’s endogenous pacemaker system and blocking electrical activation of the ventricles (***Figures 7A* *and S7B-D***). When we paced the guinea pig hearts at 2Hz, the optical action potentials from the AVNLPC and VLCM grafts (ASAP1) and the host hearts (RH237) were synchronized, indicating that the grafts are electrically coupled with the host^70^ (n = 4 of 6 animals AVNLPCs, n = 7 of 8 animals VLCMs, ***Figures 7D and 7F***). Additionally, graft-derived optical action potentials were aligned with the heart’s ECG, further confirming electrical coupling.

**Figure 7:**
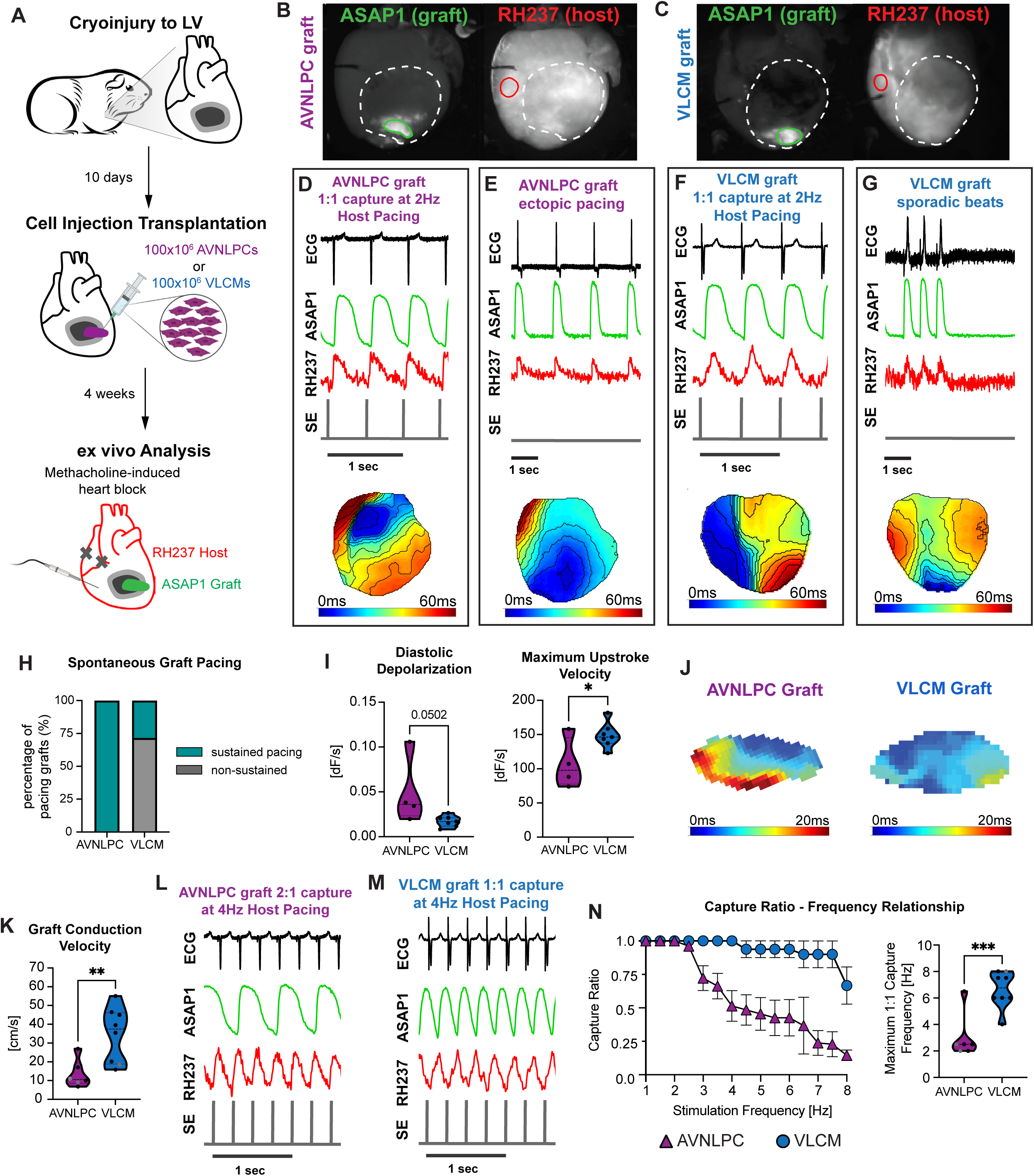
AVNLPCs exhibit AVN conduction properties in vivo. **A.** Schematic overview of the guinea pig cryoinjury model used to assess the AVNLPC conduction properties in vivo. Ten days after cryoinjury cells are injected into the border zone of the scar. Hearts are analyzed ex-vivo after 4 weeks using the Langendorff system together with a dual optical mapping setup allowing to confirm electrical integration of graft with the host heart. To control the heart rate, hearts are perfused with methacholine inhibiting the guinea pigs SAN and AVN and subsequently paced from the right ventricle using a stimulation electrode. **B, C.** Representative images of hearts containing an ASAP1-derived AVNLPCs graft B. or VLCM graft C. acquired in the ASAP1 (green) and RH237 (red) channels. The white dotted line indicates the scar area. Optical action potentials (oAP) of the AVNLPC/VLCM grafts are recorded in the region of interest (ROI) marked by the green solid line, while host oAPs are recorded in the ROI marked by the red solid line. **D, E.** Representative host ECG traces together with oAPs recorded in the AVNLPC graft (green) and in the host (red) and the signal form the stimulation electrode (grey) showing 1:1 graft-host coupling at 2Hz pacing D. and sustained spontaneous pacing of the AVNLPC graft when external pacing is terminated (flat grey line) E., in methacholine perfused hearts. The bottom of each panel shows the voltage activation map of the heart derived from the RH237 signal indicating the origin of pacing. **F, G.** Representative host ECG traces together with oAPs recorded in the VLCM graft (green) and in the host (red) and the signal form the stimulation electrode (grey) showing 1:1 graft-host coupling at 2Hz pacing F. and non-sustained spontaneous pacing of the VLCM graft when external pacing is terminated (flat grey line) G., in methacholine perfused hearts. The bottom of each panel shows the voltage activation map of the heart derived from the RH237 signal indicating the origin of pacing. **H.** Summary of the proportion of AVNLPC and VLCM grafts showing sustained vs non-sustained pacing (AVNLPC grafts: n = 4, VLCM grafts: n = 7). **I.** Quantification of indicated optical action potential parameters in AVNLPC and VLCM grafts (AVNLPC grafts: n = 4, VLCM grafts: n = 7). **J.** Representative voltage activation maps of AVNLPC and VLCM grafts derived from the ASAP1 signal. **K.** Quantification of conduction velocities recorded in AVNLPC and VLCM grafts (AVNLPC grafts: n = 6, VLCM grafts: n = 8). Grafts that were not electrically coupled are indicated as grey dots. **L.** Representative host ECG traces together with oAPs recorded in the AVNLPC graft (green) and in the host (red) and the signal form the stimulation electrode (grey) showing 2:1 graft-host coupling at 4Hz pacing. **M.** Representative host ECG traces together with oAPs recorded in the VLCM graft (green) and in the host (red) and the signal form the stimulation electrode (grey) showing persistent 1:1 graft-host coupling at 4Hz pacing. **N.** Graph summarizing the capture ratios of AVNLPC and VLCM grafts at increasing pacing frequencies (left). Quantification of the pacing frequency at which 1:1 capture is lost (right) (AVNLPC grafts: n = 6, VLCM grafts: n = 8) Grafts that were not electrically coupled are indicated as grey dots. Error bars represent SEM. Statistical tests: unpaired two-sided t-test (I, K, N): *P < 0.05, **P < 0.001.

Since the AVN functions as a secondary pacemaker, we aimed to evaluate the ability of the AVNLPC grafts that were electrically coupled with the host to pace the guinea pig hearts^71^ (***Figure 7E***). When we ceased electrical pacing, all four animals with AVNLPC grafts displayed sustained ventricular ectopic beats, suggesting that the graft was pacing the ventricle. ECG recordings (change in direction) and voltage activation maps (change in the location of signal initiation) confirmed that these ectopic beats originated from the graft site. In contrast, only two out of seven animals with electrically coupled VLCM grafts displayed sustained ectopic pacing while the remaining five animals exhibited only sporadic ectopic beats (***Figures 7G and 7H***). The spontaneous action potentials of the AVNLPC grafts retained the pacemaker phenotype seen in vitro, exhibiting diastolic depolarization and slower maximum upstroke velocity compared to VLCM grafts (***Figure 7I***). As observed in vitro, there was no significant difference in APD30 or APD90 between AVNLPC and VLCM grafts (***Figures S7E***). We next used this model to assess the conduction properties of the AVNLPC grafts. Electrical coupling between the graft and host allowed us to pace the right ventricle and assess conduction within the graft using the ASAP1 signal. In cases where electrical coupling was absent, grafts were directly paced with the external electrode. When we paced the hearts/grafts at 2Hz, the AVNLPC grafts displayed conduction velocities within the reported range of human AVN conduction (13.5±7.4 cm/s)^1,72^, which were significantly slower compared to the VLCM grafts (34.5±14.7 cm/s) (***Figures 7J and 7K***). To assess whether AVNLPC grafts can block the conduction of rapid rhythms *in vivo*, we paced the hearts/grafts at increasing frequencies (***Figures 7L* *-7N*)**. The AVNLPC grafts sustained a 1:1 capture ratio up to 3.0±0.7 Hz, while the VLCM grafts maintained a 1:1 capture ratio up to 6.6±1.4 Hz. Finally, hearts were sectioned and stained for human specific KU80 to identify the human cell grafts (***Figures S7F and S7G***). AVNLPC grafts stained cTNT^+^ MSX2^+^ while VLCM grafts stained cTNT^+^ MSX2^low^ indicating that the grafts maintained their molecular identify during the four-week engraftment period. Together, these experiments show that AVNLPCs can function as an ectopic pacemaker. Most importantly, they demonstrate that AVNLPCs maintain their AVN phenotype and conduction properties upon engraftment into the heart and are more effective in blocking the conduction of fast rates compared to VLCMs.

## Discussion

Although several protocols for differentiating hPSCs into AVN-like pacemaker cells have been described^43–46^, a comprehensive functional evaluation to determine their potential for biological conduction bridge (BioCB) applications is still lacking. Here, we report an efficient BMP signaling-based approach for generating AVNLPCs from multiple hPSC lines. We show that these AVNLPCs transcriptionally and functionally mimic AVN pacemaker cells. Importantly, AVNLPCs display slow decremental conduction, and recapitulate the AVN’s crucial safety function of blocking fast atrial rates both in vitro and in vivo. These findings establish a foundation for future in vitro modelling of AVN diseases and the development of a BioCB as a potential treatment for patients with AV block.

In agreement with prior studies in animal models, we demonstrate that WNT activation using the small molecule CHIR99021 can induce the differentiation of hPSC-derived cardiac progenitors towards AVNLPCs^33,34^. Downstream signaling analysis identified BMP as a key mediator of this effect, aligning with established developmental literature^2,35^. Accordingly, activation of BMP signaling using BMP2 was equally potent in inducing AVNLPC differentiation. Of note, our experiments showed that appropriate developmental timing of WNT or BMP activation at the heart tube stage (days 5-11) is required for the efficient induction of AVNLPC differentiation.

Interestingly, we found a baseline level of NKX2-5^+^TBX3^+^ cardiomyocytes using standard ventricular cardiomyocyte differentiation protocols that could be blocked by inhibition of BMP signaling. For cell therapy applications aimed at treating myocardial infarction, where the presence of pacemaker cells would be detrimental, depletion of unwanted TBX3^+^ pacemaker cells could be highly beneficial^73,74^. Our findings suggest that incorporating a BMP inhibition step immediately following the specification of cardiac progenitors could present a viable strategy to remove pacemaker cells from VLCM differentiations and enhance the safety of VLCM-based cell therapies.

Using multiple hPSC lines we found that CHIR99021 treatment efficiently induced an AVNLPC phenotype in cardiac progenitors but reduced the overall cardiomyocyte content at the end of differentiation. In contrast BMP2 treatment efficiently induced AVNLPCs while maintaining a high cardiomyocyte content for all cell lines tested. This establishes a robust, reproducible protocol for generating AVNLPCs from hPSCs.

Four other studies reported the generation of AVC/AVN-like cardiomyocytes from hPSCs. The AVC/AVN-like cardiomyocytes reported by Du et al, Ye et al, and Li et al. were specified using a WNT-based protocol while Schmidt et al. used a BMP-based protocol^43–46^. Our differentiation approach is unique from these previous studies by utilizing embryoid bodies in suspension culture rather than adherent monolayer cultures. Embryoid body-based protocols offer the advantage of easy adaptation to bioreactors for large scale cell manufacturing^75^. The use of the HES3 NKX2-5^eGFP/wt^TBX3^tdTomato/wt^ dual reporter line allowed us to precisely quantify the proportion of AVNLPCs in all experimental conditions. To validate the protocol in additional hPSC lines, we quantified AVNLPC proportions using MSX2 immunofluorescent staining. With that we were able to show differentiation efficiencies of 60-70% for multiple hPSC lines. In contrast, previous studies either lacked quantification across multiple differentiations and cell lines or reported lower differentiation efficiencies^43–46^.

We used a single cell transcriptome-based approach to further validate the phenotype of our hPSC-derived AVNLPCs. We were able to show that AVNLPCs closely resembled the pacemaker cells of fetal AVN, expressing key AVN markers while lacking expression of working cardiomyocyte markers. These findings confirm that BMP2 treatment at the cardiac progenitor stage effectively guides specification towards the AVN lineage.

Studies of mouse, rabbit and human hearts have described the AVN as a heterogenous structure consisting of a Core AVN that is surrounded by transitional cell types, including atrial transition zone (ATZ) cells toward the right atrium and Lower Nodal Bundle (LNB) cells toward the right ventricle and His bundle^5,76–78^. Our scRNA-seq analysis of the fetal AVN recapitulated this heterogeneity. Core AVN cells were identified based on their high expression of AVN and pan-pacemaker markers. ATZ cells expressed a mix of atrial and AVN markers, consistent with prior reports describing these cells as transitioning from an atrial to AVN phenotype^78,79^. The LNB cells expressed a combination of ventricular and AVN markers, similar to the transcriptomic profile of LNB cells of the mouse heart recently characterized by Oh et al^80^. Importantly, the hPSC-derived AVNLPCs mirrored the in vivo AVN composition, containing cells resembling the core AVN, ATZ and LNB. This is the first time that these AVN cardiomyocyte subtypes have been generated from hPSCs and underscores the strength of our developmental biology-based approach in capturing native AVN features. The presence of AVN cardiomyocyte subtypes, offers the potential to refine differentiation strategies - either enhancing core AVN purity or enriching for transitional cell types to improve future BioCB and disease modeling applications.

Studies in the mouse heart showed that both first heart field (FHF) and posterior second heart field (pSHF) progenitors contribute to the development of the atrioventricular canal^81–84^. Of note, the contribution of these progenitors to the definitive AVN has not yet been fully elucidated. Our optimized AVNLPC differentiation protocol transitions through a RALDH2^−^CD235^+^ mesoderm suggesting a FHF rather than pSHF origin^29,85^. Accordingly, the addition of RA to this mesoderm significantly reduced the AVNLPC population further supporting a FHF origin. Previously published AVC/AVN hPSC protocols incorporated RA signaling activation either directly through RA supplementation, or indirectly via retinol containing media, suggesting that these AVC/AVN-like cardiomyocytes derive from a pSHF progenitor^43–46^. Notably, Schmidt et al. and Ye et al. reported high expression of the fast-conducting Cx40 (*GJA5*) in their AVC/AVN-like cardiomyocytes which is usually expressed by atrial cardiomyocytes, another pSHF derivative^44,46^. In contrast, our AVNLPCs exhibit low *GJA5* expression, aligning more with an AVN phenotype. Future studies are needed to further investigate the molecular and functional differences between FHF and pSHF-derived AVC/AVN-like cardiomyocytes and how this might impact their application towards disease modelling and cell therapies.

The timing and mechanisms governing the specification of AVC cardiomyocytes into AVN cardiomyocytes in the human heart remain unknown. Moreover, there are currently no specific markers to distinguish AVC cardiomyocytes from AVN cardiomyocytes of the definitive core AVN. As a result, it is challenging to determine whether our hPSC-derived AVNLPCs represent the core AVN population or resemble earlier AVC cardiomyocytes, given the significant overlap in AVC and AVN gene expression. However, in the mouse heart the AVN is established by E17^41,86^, which corresponds to approximately gestation week 10 of human development^87–89^. Based on this developmental timeline, the Core AVN cluster identified in our gestation week 18 fetal AVN sample represents AVN cardiomyocytes. Spearman correlation analysis demonstrated similarity between our AVNLPCs with this fetal AVN cardiomyocyte cluster suggesting an AVN phenotype. Additionally, our AVNLPCs exhibited a high degree of alignment with adult AVN DEGs, further supporting their AVN-like state. In addition to differentiating into AVN cardiomyocytes, AVC cardiomyocytes differentiate into atrial and ventricular chamber cardiomyocytes^37,38^. Our in vivo experiments demonstrated that AVNLPC grafts maintained a stable AVN-like phenotype for up to four weeks, as indicated by the expression of AVN marker MSX2 and functional properties including slow conduction velocity and the ability to block conduction of fast pacing rates. Based on this we refer to our cells as AVN-like pacemaker cells (AVNLPCs).

The AVN is the secondary pacemaker of the heart, and as such, AVNLPCs express the pacemaker marker *TBX3*. Patch clamp and optical action potential recordings showed AVNLPCs display typical pacemaker action potentials with faster diastolic depolarization and larger underlying pacemaker funny currents (I_f_) compared to VLCMs. Under physiological conditions the heartbeat is initiated by the primary pacemaker the SAN and the AVN serves as a conduction bridge, transmitting electrical impulses from the atria to the ventricles. To thoroughly investigate the conduction properties of our AVNLPCs, we integrated the cells into engineered heart tissues (EHTs) containing cardiac fibroblasts to closely mimic genuine cardiac tissue architecture. These tissues compact around flexible poles, generating mechanical force that is known to promote cardiomyocyte maturation^32,90^ – a feature absent from other published approaches that relied on models like monolayers, fused spheroids or 3D assembloids^43–46^.

Utilizing this physiologically relevant platform we found that AVNLPCs integrated into EHTs (referred to here as engineered AVN tissues (eAVNTs)) exhibit conduction velocities of ∼4 cm/s comparable to the human AVN^1^. Accordingly, eAVNTs exhibit significantly slower conduction velocities than engineered ventricular tissues (eVTs) containing VLCMs which is in agreement with previous reports of hPSC-derived AVC/AVN-like cells^43–46^.

What is more, we demonstrate that AVNLPCs recapitulate hallmark features of AVN electrophysiology. When eAVNTs were subjected to incremental pacing to simulate atrial fibrillation, they blocked conduction of impulses above 3.3 Hz, effectively filtering high-frequency signals and conducting only every other beat at higher rates. Analysis of the underlying mechanism revealed that eAVNTs display the Wenckebach phenomenon in response to fast pacing rates. Further, when paced with premature stimuli, eAVNTs demonstrated decremental conduction. The replication of these complex AVN properties in hPSC-derived cells is remarkable and provides strong evidence that we generated bonafide AVN cells. These findings go well beyond previous reports that only described slower conduction in AVC/AVN-like cells^43–46^. It is possible that the use of EHTs that more closely mimic cardiac tissue contributed to the advanced functional properties we observed. Notably, these functional properties of the AVNLPCs are critical for the development of a BioCB that needs to replicate the AVNs role as a gatekeeper. Specifically, the ability to progressively slow signal propagation in response to rapid atrial rates, thereby preventing tachyarrhythmias from being transmitted to the ventricles where they would be life-threatening. What is more, we show that eVTs composed of VLCMs were less effective in blocking high-frequency impulses and lacked these crucial AVN conduction properties. This underscores the importance of generating the correct cardiomyocyte subtype for future therapeutic application as a BioCB.

To enable future BioCB applications, it is important to confirm that AVNLPCs maintain their AVN-like phenotype following engraftment in vivo. Here, we used state-of- the-art genetically encoded voltage sensors and voltage sensitive dyes that allowed us to assess electrical integration of the graft with the host and graft conduction properties. We demonstrate, for the first time, that hPSC-derived AVNLPCs retain their AVN-like functional properties when transplanted into a cryoinjury guinea pig model, marking a key advancement over prior studies focused solely on in vitro characterization^43–46^. We show that AVNLPCs electrically couple with host cardiomyocytes and pace the host heart upon pharmacological SAN and AVN block, demonstrating the ability to function as a backup pacemaker like the genuine AVN. Furthermore, AVNLPCs maintained their conduction properties for up to four weeks post-transplantation. Although conduction velocity in vivo was faster than in vitro (13.5±7.4 cm/s vs. 3.817±1.5cm/s), which likely reflects graft maturation over time, the conduction velocity of the AVNLPCs remained within the physiological range of the human AVN. Importantly, AVNLPC grafts also demonstrated the ability to block simulated atrial arrhythmias at pacing rates above 3.0 Hz (180 bpm), in contrast to VLCM grafts that only blocked rates above 6.6 Hz (396 bpm). Conduction of a rate of 6 Hz to the ventricles would be life-threatening in humans, again, underscoring the importance of generating the appropriate cell type for the therapeutic application in question - in this case AVNLPCs and not VLCMs, for safe and effective future BioCB therapies.

### Limitations

The AVN is a heterogenous tissue containing core AVN pacemaker cells as well as transitional cell types with mixed phenotypes at the border zones towards both the right atrium and His Bundle. scRNA-seq analysis of the AVNLPCs revealed that they represent a mix of core AVN pacemaker cells, ATZ cells and LNB cells. Future studies will focus on identifying the signaling pathways that specifically drive the differentiation of each of these cardiomyocyte subtypes. Although AVNLPCs maintain their function in vivo, their ability to serve as a BioCB was not fully determined. To properly test the ability of AVNLPCs to function as a BioCB it will be required to achieve engraftment spanning from the atria to the ventricles. This was challenging to accomplish in a small animal model and attempts to suture the engineered heart tissues into the AV-groove of the guinea pig heart failed due to the lack of electrical integration of the grafts with the host hearts. Larger animal models that allow for better surgical manipulation of the heart will have to be employed to deliver the cells into the AVN region to address this question. Finally, like every hPSC-derived product AVNLPCs are immature and do not represent AVN pacemaker cells of the adult heart. However, we were able to show striking functional differences between AVNLPCs and VLCMs demonstrating that these cells are a valuable tool for in vitro modeling of AVN diseases and the development of a BioCB as improved future cell therapy-based treatment for AV-block patients.

## Methods

### KEY RESOURCES TABLES

**Table S1:** Marker genes used to identify cell types.

| Cell Type | Marker Genes |
| --- | --- |
| Cardiomyocytes (general) | TNNT2, TNNI3, ACTN2, MYH6 |
| Atrial Cardiomyocytes | NKX2-5, NR2F2, NPPA, GJA5, SCN5A |
| Atrial Transition Zone | NKX2-5, TBX3, HCN4, GJC1, NR2F2, NPPA, GJA5, SCN5A |
| AVN Core/Compact Cardiomyocytes | NKX2-5, TBX3, HCN4, GJC1, MSX2, TBX2, BMP2, RSPO3 |
| His Bundle | NKX2-5, TBX3, HCN4, GJC1, MSX2, TBX2, BMP2, RSPO3, SCN5A, GJA5 |
| SAN Cardiomyocytes | TBX3, HCN4, GJC1, TBX18, SHOX2, ISL1, CD34, MYH11 |
| Ventricular Cardiomyocytes | NKX2-5, MYL2, IRX4, GJA1, SCN5A |
| Lower Nodal Bundle | NKX2-5, TBX3, HCN4, GJC1, MYL2, IRX4, GJA1, SCN5A |
| Endocardial | NFATC1, NRG1, CDH11, NPR3, FABP4, PLVAP |
| Endothelial | KDR, CDH5, PECAM1 |
| Epicardial | UPK3B, ALDH1A2, WT1, TBX18 |
| Epithelial | KRT19, CLDN4, ALCAM, EPCAM, CDH1 |
| Fibroblasts | PDGFRA, TCF21, THY1, VIM, DDR2 |
| Macrophages | CSF1R, CD74, AIF1, CD163, CD68 |
| Neuronal | RBFOX3, PAX6, NES, GFAP, TUBB, SOX2, ASCL1 |
| Proliferating | KI67, CDK1, PIF1, FOXM1, TOP2A |
| Smooth Muscle | MYH11, ACTA2, TAGLN, CALD1, MYOCD |

**Table S6:**
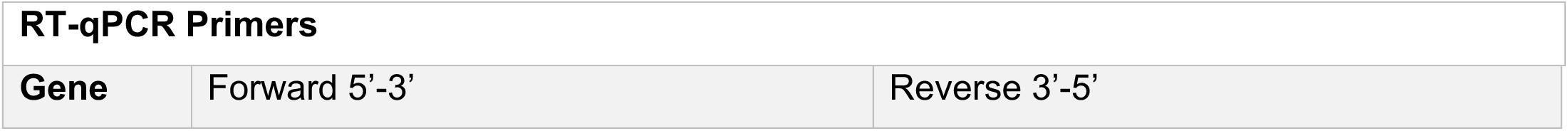
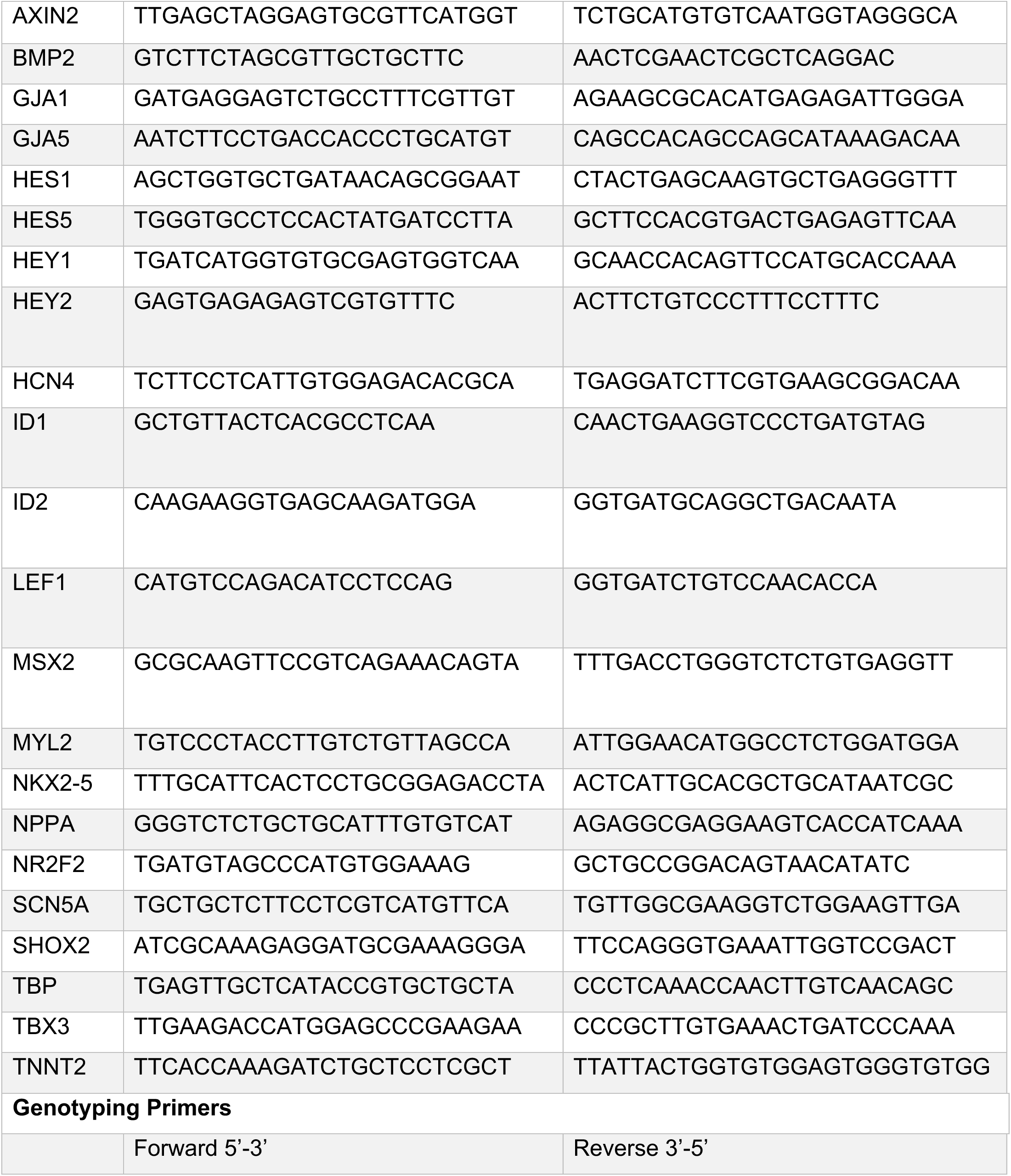
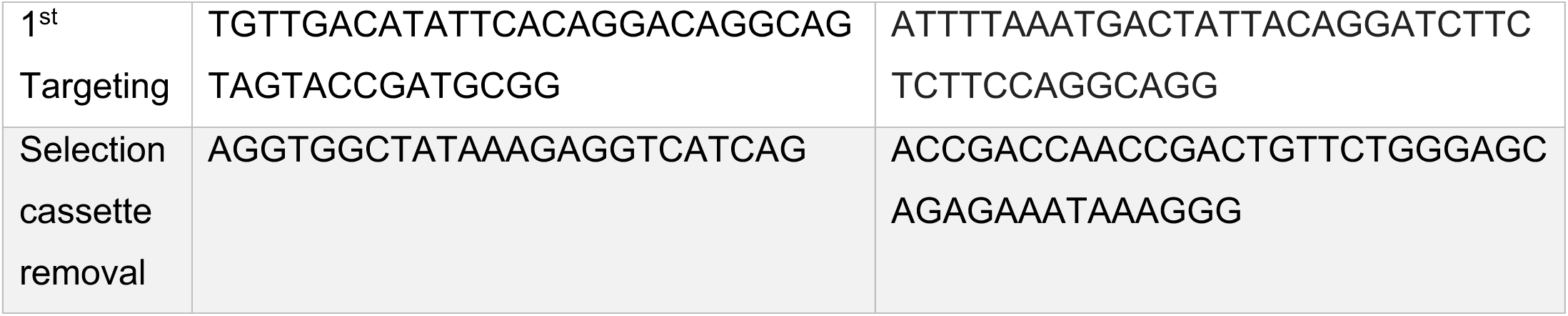
Primer List.

**Table S7:** Animal Reagents.

| Reagent | Manufacturer |
| --- | --- |
| Ketamine hydrochloride (Narketan) | Vetoquinol, DIN02374994 |
| Xylazine (Rompun) | Elanco, DIN02169592 |
| Meloxicam (Metcam) | Boehringer Ingelheim Animal Health, 02240463 |
| Buprenorphine-slow release (Vetergesic Mulidose) | CEVA Animal Health, DIN02342510 |
| Cyclosporin A/Sandimmune IV | Novartis, 00593257 |
| Methylprednisolone (Solu-Medrol) | Pfizer, DIN02367971 |

**Table S8:** Chemical Reagents.

| Reagent | Manufacturer |
| --- | --- |
| Calcium chloride – CaCl<sub>2</sub> | Fisher (Fluka), 15641930 |
| D-Glucose – dextrose anhydrous powder | Sigma, G8270 |
| HEPES | Bioshop, 7365-45-9 |
| Magnesium ATP – MgATP | Sigma A9187 |
| Magnesium chloride – MgCl<sub>2</sub> | Fisher (Fluka), 60047233 |
| Magnesium sulfate – MgSO<sub>4</sub> | Sigma, M1880 |
| Potassium aspartate | Sigma, A6558 |
| Potassium chloride – KCl | Sigma, P4504 |
| Potassium phosphate monobasic – KH<sub>2</sub>PO<sub>4</sub> | Sigma, P5655 |
| Sodium bicarbonate – NaHCO<sub>3</sub> | Sigma, S5761 |
| Sodium chloride – NaCl | Sigma, S0876 |
| Sodium hydroxide – NaOH | Sigma, 221465 |
| Sodium phosphate monobasic – NaH<sub>2</sub>PO<sub>4</sub> | Sigma, S0751 |
| Sodium pyruvate – C<sub>3</sub>H<sub>3</sub>NaO<sub>3</sub> | Sigma, P2256 |

**Table S9:** Primary antibodies used for flow cytometry and cell sorting.

| Antibody | Manufacturer | Concentration |
| --- | --- | --- |
| anti-PDGFRa-PE | BD, 556002 | 6:100 |
| anti-CD56-APC | BD, 555518 | 1:100 |
| anti-CD235a-APC | BD, 551336 | 1:100 |
| anti-CD172a/b-PE-Cy7 (SIRPa) | Biolegend, 323808 | 1:1000 |
| anti-CD90-APC | BD, 559869 | 1:300 |
| anti-Cardiac Troponin T (cTNT) | ThermoFisher, MA5-12960 | 1:2000 |
| anti-MLC2v | Abcam, ab79935 | 1:1000 |
| anti-OCT3/4-PE | BD, 561556 | 1:300 |
| anti-SOX2-PerCP-Cy5.5 | BD, 561506 | 1:100 |

**Table S10:** Secondary antibodies used to detect primary antibodies for flow cytometry.

| Antibody | Manufacturer | Concentration |
| --- | --- | --- |
| goat anti-mouse IgG-A647 | BD, 550826 | 1:500 |
| donkey anti-rabbit IgG-PE | Jackson ImmunoResearch, 711-116-152 | 1:500 |

**Table S11:**
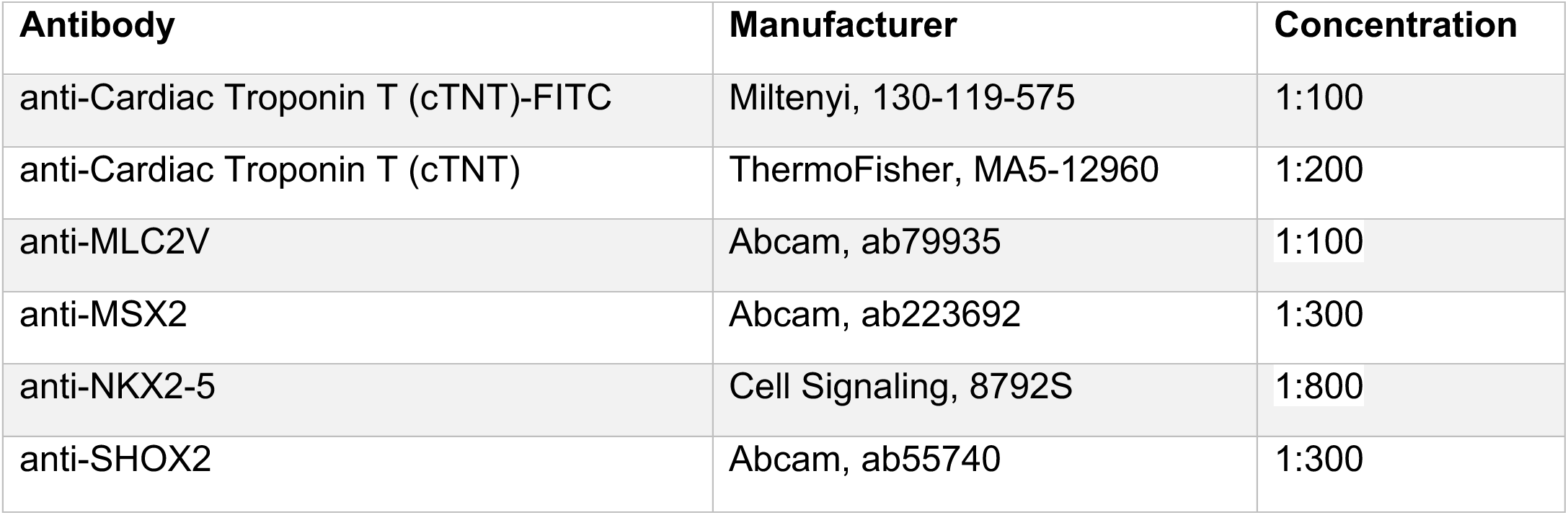
Primary antibodies used for monolayer and section staining.

| Antibody | Manufacturer | Concentration |
| --- | --- | --- |
| anti-Cardiac Troponin T (cTNT)-FITC | Miltenyi, 130-119-575 | 1:100 |
| anti-Cardiac Troponin T (cTNT) | ThermoFisher, MA5-12960 | 1:200 |
| anti-MLC2V | Abcam, ab79935 | 1:100 |
| anti-MSX2 | Abcam, ab223692 | 1:300 |
| anti-NKX2-5 | Cell Signaling, 8792S | 1:800 |
| anti-SHOX2 | Abcam, ab55740 | 1:300 |

**Table S12:** Secondary antibodies used for monolayer and section staining.

| Antibody | Manufacturer | Concentration |
| --- | --- | --- |
| Donkey anti-rabbit IgG-A647 | ThermoFisher, A31573 | 1:500 |
| Donkey anti-rabbit IgG-A555 | ThermoFisher, A31572 | 1:500 |
| Donkey anti-mouse IgG-A647 | ThermoFisher, A31571 | 1:500 |
| Donkey anti-mouse IgG-A555 | ThermoFisher, A31570 | 1:500 |

### CONTACT FOR REAGENT AND RESOURCE SHARING

Further information and requests for resources and reagents should be directed to and will be fulfilled by the Lead Contact, Stephanie Protze.

### EXPERIMENTAL MODEL AND SUBJECT DETAILS

#### Generation and maintenance of human ESC lines

All human pluripotent stem cell (hPSC) research conducted in this study was approved by the Stem Cell Oversight Committee of the Canadian Institutes of Health Research and by UHN’s Research Ethics Board. The HES3-Nkx2-5^egfp/w^ TBX3^tdtom/w^ reporter cell line (karyotype: 46, XX)^47^ was generated as detailed in the methods section below. The HES2 cell line (karyotype: 46, XX)^57^ was purchased from WiCell. The ESI-017 ASAP1 cell line (karyotype: 46, XX)^56^ was generously provided by Dr. Michael Laflamme.

#### Generation of the HES3-Nkx2-5^egfp/w^ TBX3^tdtomato/w^ stem cell line

The Hes3 NKX2-5^eGFP/wt^ human embryonic stem cell (hESC) line was a kind gift from Prof. E. Stanley (Monash University, Victoria, AU)^47^. The introduction of the tandem dimer tomato (tdTomato) fluorescence reporter gene into the TBX3 locus of this cell line was performed by the Centre for Commercialization of Regenerative Medicine, Toronto. TALEN pairs were designed and tested for cutting efficiency in collaboration with Cellectis bioresearch, France. The left and right DNA binding TALEN sequences were cloned in two separate backbones by Cellectis bioresearch and sequence verified. TALENS were selected based on cutting efficiency and to target the endogenous start codon (ATG) of TBX3 to incorporate the tdTomato fluorescence gene (***Figure S1A***). The targeting vector comprised of: 1kb homology arms on each side of the endogenous start codon of TBX3, coding sequence of tdTomato, a polyadenylation signal followed by a FRT flanked Neomycin selection cassette under a ubiquitous human phosphoglycerate kinase promoter (hPGK). The sequence “AGCCTCTCCATGAGAGATCCG” was removed from targeting vector to prevent TALEN binding and cutting of the targeting vector. The targeting vector was gene synthesized by GENEART (ThermoFisher Scientific), endotoxin free maxi prepped and sequence verified.

For transgene delivery, Hes3 NKX2-5^eGFP/wt^ cells were trypsinized with TrypL (ThermoFisher, 12605010) into a single cell suspension and 800,000 cells were electroporated with 2.5 ug each of the two TALEN plasmids along with 2 ug of targeting vector using the DN100 program of an Amaxa 4D Nucleofector and P3 primary cell solution (Lonza, V4XP-3034). The electroporated cells were plated onto drug resistant Irradiated DR4 Mouse embryonic fibroblasts (ThermoFisher, A34966) in three 10 cm cell culture dishes and successfully targeted hESCs were selected using G418 treatment (ThermoFisher, 10131035).

Clones with heterozygous edited TBX3 gene locus were identified by PCR screening and confirmed using Sanger sequencing (1^st^ targeting primers, ***Table S6***). To remove the neomycin selection cassette, a successfully edited Hes3 NKX2-5^eGFP/wt^ clone was transfected with an in vitro transcribed mouse codon optimized FLP recombinase mRNA (Miltenyi Biotec, 130-106-769). Excision of the neomycin selection cassette was confirmed via PCR and sanger sequencing (Selection cassette removal primers, ***Table S6***).

The final NKX2-5^eGFP/wt^ TBX3^tdTomato/wt^ hES clones (#5 and #7) were subjected to Mycoplasma testing (negative). Normal karyotype (46, XX) was confirmed by Standard G banding karyotyping by the Institute of Human Genetics, University Clinic Jena, Germany (***Figure S1C***). Pluripotency was confirmed via flow cytometry using anti-Oct3/4-PE (BD, 561556), anti-Sox2-PerCP-Cy5.5 (BD, 561506) (***Figure S1D***).

#### Human pluripotent stem cell maintenance

Human pluripotent stem cell lines were cultured as described previously^91^. Briefly, cells were maintained as monolayers on a 0.1% gelatin and 0.1% Matrigel (FisherScientific, CB40230) matrix with irradiated mouse embryonic fibroblasts in DMEMF/12 (ThermoFisher, MT-10-092-CV) supplemented with 20% KnockOut serum replacement (ThermoFisher, 10828028), 2mM L-glutamine (ThermoFisher, 25030081), 1x non-essential amino acids (ThermoFisher, 11140050), 55mM β-Mercaptoethanol (ThermoFisher, 21985023) and 1% penicillin/streptomycin (ThermoFisher, 15070-063), and 20ng/ml recombinant human (rh)-bFGF (Biotechne, 233-FB/CF). All hPSC lines had normal karyotypes and tested negative for mycoplasma.

#### Use of human fetal tissue

Human fetal heart sample from gestation weeks 19 was provided by the Toronto Mount Sinai Hospital Research Centre for Women’s and Infants’ Health Biobank. The use of human fetal tissue in this study was approved by the Research Ethics Boards of both Mount Sinai Hospital, Toronto, and the UHN, Toronto. Samples were obtained from voluntary pregnancy terminations. After patients signed the standard consent form for the procedure, they were approached by the Mount Sinai Hospital Research Centre for Women’s and Infants’ Health BioBank to request consent for the donation of fetal tissue. Donors were fully informed about the intended research use of the samples and did not receive compensation for their donation. Fetuses with chromosomal abnormalities, cardiovascular defects, or from mothers with congenital heart defects, arrhythmias, diabetes (type 1, type 2, or gestational), or Systemic Lupus Erythematosus were excluded from the study to ensure the inclusion of healthy heart samples. The fetal sex was not specified for this study and not included in the health information shared with the study team.

#### Use of Animals

##### Guinea Pig use in experimental studies

Male Hartley guinea pigs (650–700 g, Charles River) were used, with all procedures approved by the Animal Use and Care Committee at UHN. Surgeries were performed under ketamine (35mg/kg, Vetoquinol, DIN02374994) and xylazine (2mg/kg, Elanco, DIN02169592) induction and isoflurane maintenance, with orotracheal intubation and mechanical ventilation. Meloxicam (0.2 mg/kg, Boehringer Ingelheim Animal Health, 02240463) and Buprenorphine-slow release (0.7-1.2 mg/kg, CEVA Animal Health, DIN02342510) were administered for pain management. Animals were placed on a 38°C water blanket to maintain body temperature, O2 saturation and heart rate were monitored using a toe probe. The surgical site was prepared with betadine, and ophthalmic ointment was applied to prevent corneal drying. Postoperatively, animals received subcutaneous saline (5 mL) and were monitored in a warm environment until full recovery. They were assessed daily for the first week, then biweekly, with immunosuppressed animals weighed and examined daily. Veterinary services were consulted if signs of severe distress, infection, or >20% weight loss were observed.

### METHODS DETAILS

#### Directed Differentiation of Human ESC Lines

To differentiate hPSCs into cardiomyocytes, we employed our previously reported embryoid body (EB)-based protocol^29,32,92^. For all differentiation steps, the base media consisted of either 100% StemPro-34 media (ThermoFisher, 10639011) or 20% StemPro-34 media diluted with IMDM (ThermoFisher, 12440061). Base media was supplemented with 1% penicillin/streptomycin, 2 mM L-glutamine, 50 μg/mL ascorbic acid (MilliporeSigma, A4544), 75 μg/mL transferrin (MilliporeSigma, 10652202001), and 50 μg/mL monothioglycerol (MilliporeSigma, M6145). hPSC colonies were grown to ∼80% confluence and dissociated into single cells using TrypLE (ThermoFisher, 12605010). Cells were aggregated for 18 hours on an orbital shaker at 70rpm (ESI-017 ASAP1) or 80rpm (HES3, HES2) in 6cm low-attachment dishes at a concentration of 1×10^6^ cells/mL. Cells were suspended in 20% StemPro-34 base media supplemented with 0.01x ITS-X (ThermoFisher, 51500056), 1ng/mL rhBMP4 (Biotechne, 314-BP/CF), 10μM ROCK inhibitor Y27632 (Selleckchem, S1049), and 250U/ml DNase I (MilliporeSigma, 260913). EBs were incubated in a low oxygen environment (5% CO_2_, 5% O_2_, 90% N_2_) until day 11 and then in a 5% CO_2_ ambient air environment thereafter. The EBs were kept in tissue culture plastic pre-coated with 5% (w/v) poly-HEMA (MilliporeSigma, P3932) throughout the differentiation to maintain suspension cultures.

#### Generation of ventricular-like cardiomyocytes (VLCMs)

On day 1 of differentiation, EBs were transferred to ventricular mesoderm induction media consisting of cardiac base media: 100% StemPro-34 supplemented with 1 μM CHIR99021 (Biotechne, 4423), 2.5 ng/mL rhbFGF, 10 ng/mL rhBMP4, and 8 ng/mL rhActivin A (Biotechne, 338-AC/CF). On day 3, EBs were washed with IMDM and suspended in cardiac induction media containing 100% StemPro-34 with 1 μM IWP2 (Biotechne, 3533) and 10 ng/mL rhVEGF (Biotechne, 293-VE/CF). On day 5, the media was replaced with cardiac maintenance media: 20% StemPro-34 and 5 ng/mL rhVEGF. Starting from day 11, EBs were cultured in maintenance media without rhVEGF. Media changes were performed every 3–4 days.

#### Generation of atrial-like cardiomyocytes (ALCMs)

At day 1 of differentiation the EBs were transferred to atrial mesoderm induction media consisting of: cardiac base media of 100% StemPro-34 with 1μM CHIR99021, 2.5ng/ml rhbFGF, 3ng/ml rhBMP4, and 2ng/ml rhActivinA. At day 3 the EBs were washed with IMDM and suspended in cardiac induction media (as described for VLCMs) with the addition of 0.5uM retinoic acid (MilliporeSigma, R2625). From day 5 onward, media changes were performed as described for VLCMs.

#### Generation of sinoatrial node-like pacemaker cells (SANLPCs)

On day 1 of differentiation, embryoid bodies (EBs) were transferred to SAN pacemaker mesoderm induction media composed of cardiac base media (100% StemPro-34) supplemented with 1 μM CHIR99021, 2.5 ng/mL rhbFGF, 3 ng/mL rhBMP4, and 2 ng/mL rhActivin A. On day 3, EBs were washed with IMDM and transitioned to SAN pacemaker induction media containing 0.5 μM IWP2, 10 ng/mL rhVEGF, 2.5 ng/mL rhBMP4, 1.5 μM SB-431542 (MilliporeSigma, S4317), and 0.125 μM retinoic acid. On day 4, 1000 nM PD173074 (Biotechne, 3044) was added to the media. Starting on day 6, cells were cultured in base media consisting of 100% StemPro-34 supplemented with 5 ng/mL rhVEGF. From day 8 onward cells were cultured in cardiac maintenance media and media changes were performed as described for VLCMs.

#### Generation of atrioventricular node-like pacemaker cells (AVNLPCs)

At day 1 of differentiation the EBs were transferred to AVN pacemaker mesoderm induction media consisting of: cardiac base media of 100% StemPro-34 with 1μM CHIR99021, 2.5ng/ml rhbFGF, and 5ng/ml rhBMP4 + 4ng/ml rhActivinA (HES3) or 5ng/ml rhBMP4 + 2ng/ml rhActivinA (HES2, ESI-017 ASAP1). At day 3 the EBs were washed with IMDM and suspended in cardiac induction media (as described for VLCMs). For BMP2-induced AVNLPCs: 200ng/mL rhBMP2 (Biotechne, 355-BM) was added to the cardiac maintenance media at either day 5 (HES3) or day 8 (HES2, ESI-017 ASAP1) of differentiation. At day 8 (HES3) or 11 (HES2, ESI-017 ASAP1) of differentiation and thereafter, media was changed as described for VLCMs. For CHIR-induced AVNLPCs: 12μM CHIR99021 was added to the cardiac maintenance media at either day 8 (HES3) or day 11 (HES2 and ESI-017 ASAP1) of differentiation in cardiac maintenance media. At day 11 (HES3) or 15 (HES2, ESI-017 ASAP1) of differentiation and thereafter, media was changed as described for VLCMs.

Aggregation experiments: To examine the effects of BMP and WNT signaling on AVNLPC development, embryoid bodies (EBs) were dissociated into single cells on either day 5 or day 8. Day 5 EBs were dissociated using TrypLE, while day 8 EBs were treated with Collagenase Type II (1 mg/mL, Worthington, 4176) in Hank’s buffer (NaCl, 136 mM; NaHCO₃, 4.16 mM; Na₃PO₄, 0.34 mM; KCl, 5.36 mM; KH₂PO₄, 0.44 mM; dextrose, 5.55 mM; HEPES, 5 mM (Key Resources ***Table S8)***). To promote aggregate formation, cells were cultured at high density (75,000 cells/well) in 96-well ultra-low attachment plates. The generated aggregates were cultured in cardiac induction media supplemented with 10μM ROCK inhibitor and the indicated concentrations and combinations of rhBMP2, CHIR99021, and LDN193189 (Biotechne, 6053). After 4 days the aggregates were transferred to 24-well plates pre-coated with 5% (w/v) poly-HEMA and cultured in cardiac maintenance media as described above.

#### Cardiomyocyte cryopreservation

To enhance engraftment and cell survival of transplanted cardiomyocytes in vivo, EBs were subjected to heat shock and a pro-survival cocktail before cryopreservation as previously described^93^. The heat shock process involved a 1 hour incubation at 42 °C, followed by a 3–5-hour incubation at 37 °C in cardiac maintenance media supplemented with 100 nM insulin-like growth factor-1 (IGF-1) (Biotechne, 291-G1). and 200 nM cyclosporine A/Sandimmune IV (Novartis, 00593257). EBs were dissociated overnight a RT with Collagenase Type II in Hank’s buffer as described above and then resuspended in CryoStor CS10 solution (StemCell Technologies, 100-1061) for preservation using a Controlled-Rate Freezer (Bio-Cool), as detailed in previous reports^73^.

#### RNA Isolation and Quantitative Real-Time PCR

Total RNA was extracted from hPSC-derived cardiomyocytes using the RNAqueous-Micro Total RNA Isolation Kit (ThermoFisher, AM1931), including RNase-free DNase treatment. Complementary DNA (cDNA) synthesis was performed using Superscript III Reverse Transcriptase (ThermoFisher, 18080044) with oligo(dT) primers and random hexamers. Quantitative reverse transcription PCR (RT-qPCR) was conducted using the QuantiNova SYBR Green PCR Kit (Qiagen, 208057) on a QuantStudio5 RT-qPCR machine (ThermoFisher), following the manufacturer’s instructions. All reactions were prepared in duplicate. As previously described, a tenfold serial dilution of human genomic DNA standards (30 ng/mL to 3.0 pg/mL) was included in each experiment to verify PCR efficiency and calculate the relative copy number of each target gene normalized to the housekeeping gene TBP^48^. Primer sequences are detailed in ***Table S6***. To facilitate visualization and comparison of expression patterns, the data were log2-transformed and displayed as a heatmap.

#### Flow Cytometry and Cell Sorting

EBs at Day 1-8 of differentiation were dissociated using TrypLE for 3-6 minutes at 37°C. Day 10-50 EBs were dissociated using Collagenase type II in Hank’s buffer as detailed above for 16 hours at room temperature. If necessary, this was followed by further dissociation with TrypLE. For surface marker staining, cells were incubated with antibodies for 30 minutes at 4°C in PBS (Corning, MT21030CM) containing 5% Fetal Bovine Serum (FBS) (FisherScientific, SH3039603) and 0.02% sodium azide (Sigma, S2002). Primary antibodies and their concentrations are detailed in Key Resources ***Table S9***. For intracellular staining, cells were fixed with 4% paraformaldehyde (PFA) (Cedarlane, 15710-S) for 10 minutes at 4°C. Staining was performed in PBS containing 0.3% bovine serum albumin (BSA, Sigma, A1470) and 0.3% Triton X-100 (Sigma, X100). Cells were incubated with primary antibodies overnight at 4°C, followed by a 30-minute incubation with secondary antibodies at 4°C. Secondary antibodies and their concentrations can be found in Key Resources ***Table S10***. After staining, cells were analyzed using either a LSRFortessa (BD) or a Cytoflex (Beckman Coulter) flow cytometer. Flow cytometric analysis was conducted using FlowJo software (BD).

The proportion of NKX2-5^+^TBX3^+^ cardiomyocytes (AVNLPCs) out of total cells was calculated by multiplying the proportion of SIRPA^+^CD90^−^ cardiomyocytes with the proportion of NKX2-5^+^TBX3^+^ cells gated out of the SIRPA^+^CD90^−^ gate.

For fluorescence-activated cell sorting, cells were suspended in IMDM with 0.5% fetal calf serum (FCS) and sorted at a concentration of 10 million cells/mL using a FACS Aria sorter (BD) at the SickKids-UHN Flow Cytometry Facility. Magnetic-activated cell sorting (MACS) was performed using the PSC-Derived Cardiomyocyte Isolation Kit (Miltenyi, 130-110-188) according to the manufacturer’s protocol.

#### Immunohistochemistry

##### Monolayer staining

Day 25 EBs were dissociated as described above and then cardiomyocytes were plated onto Matrigel (25% v/v, FisherScientific, CB40230) coated 12mm coverslips (FisherScientific, 22293232P) and cultured in cardiac maintenance media supplemented with 10μM ROCK inhibitor Y27632 (Selleckchem, S1049). After 48 hours, cells were transferred to cardiac maintenance media without ROCK inhibitor and then cultured for another 2-4 days until confluent. Monolayers were fixed with 4% PFA for 10 minutes at 4°C and then permeabilized in PBS containing 0.3% Triton X-100 and 200 mM glycerin (Sigma, G2289) for 20 minutes at room temperature (RT). Following permeabilization, samples were blocked using blocking buffer (PBS containing 10% donkey serum, 2% BSA, and 0.3% Triton X-100) for 30 minutes at RT. Primary antibodies (Key Resources ***Table S11***) were diluted in staining buffer (PBS containing 1% normal donkey serum (Jackson Cerdarlane, 017-000-121) and 0.3% Triton X-100), applied to the samples and incubated overnight at 4°C. The samples were then washed three times with wash buffer (PBS containing 0.1% BSA and 0.1% Triton X-100). Secondary antibodies (Key Resources ***Table S12***) were applied in staining buffer along with 10 µg/mL Hoechst (FisherScientific, H3570) for 1 hour at RT. Afterwards, the samples were washed three times with wash buffer, and once with distilled water before mounting with ProLong Diamond Antifade Mounting Media (FisherScientific, P36965).

##### Tissue section staining

Engineered heart tissues were fixed in 4% formalin (Sigma, F1635-1L) for 30 minutes at room temperature and embedded in 1% agarose (Froggabio, A87-500G). Guinea pig hearts were fixed overnight at 4°C in 4% formalin. Following fixation, all tissue samples were transferred to 70% ethanol and sent to the UHN Pathology Research Program for paraffin embedding and sectioning into 5 µm-thick slices. Masson’s Trichrome staining was performed by the UHN Pathology Research Program. For immunofluorescence staining, tissue sections were rehydrated using xylene and a graded ethanol series, followed by heat-induced antigen retrieval in citrate buffer (pH 6). Blocking was performed in PBS containing 10% donkey serum and 0.1% Triton X-100. The sections were then incubated with primary antibodies (Key Resources ***Table S11***) overnight at 4°C. After three PBS washes, secondary antibodies (Key Resources ***Table S12***) and 10 µg/mL Hoechst were applied in PBS with 0.1% Tween for 1 hour at room temperature. Finally, the samples were washed three times with PBS, and once with distilled water, and mounted using ProLong Diamond Antifade Mounting Media.

##### Imaging

All samples were imaged at the Advanced Optical Microscopy Facility at UHN. Monolayers were imaged using the Nikon A1R confocal microscope with 20x/0.75 NA and 40x/0.85 NA objectives, along with the NIS-Elements software. Section stains were imaged using the Leica SP8 with a 20x/0.75 NA objective and controlled via the LAS X software. Whole-slide images were captured using the ScanScope AT2 brightfield scanner (Aperio) with a 20x objective lens. Image processing and preparation were carried out using ImageJ (FIJI).

#### Single Cell Patch Clamp

To perform electrophysiological characterization using patch clamp recordings specific cell populations were isolated via FACS as described. Cells were resuspended at a concentration 1.25–5 × 10^6^ cells/ml in cardiac maintenance media (without VEGF) containing 10μM ROCK inhibitor Y27632 (Selleckchem, S1049) and filtered through a 70 μm filter. 40 μL drops of this suspension were plated onto 3 × 5 mm Matrigel-coated (10% v/v) glass coverslips in 30 mm dishes and incubated for 16-18 hours to facilitate cell attachment. Then, dishes were flooded with 2 mL of fresh media. Media was changed every four days, and recordings were performed 5–14 days post-plating.

Action potentials and membrane currents were recorded using standard patch clamp techniques in current- and voltage-clamp modes (Axopatch 200B, Molecular Devices). Signals were acquired at 5 kHz (DigiData, Molecular Devices) and analyzed with pCLAMP software (Molecular Devices). Borosilicate glass microelectrodes with tip resistances of 2–5 MΩ were used, with series resistance compensated by ∼70%.

Spontaneous action potentials and funny current (I_f_) were recorded at 37°C using the perforated patch method (nystatin). The bath solution contained (mM): 124 NaCl, 5.4 KCl, 1.2 CaCl₂, 1 MgCl₂, 5 glucose, and 10 HEPES (pH 7.4, NaOH) (Key Resources ***Table S8***). The pipette solution contained (mM): 120 potassium aspertate, 20 KCl, 5 NaCl, 1 MgCl₂, 5mM MgATP, and 10 HEPES (pH 7.2, KOH) (Key Resources ***Table S8***). I_f_ was elicited by 3-second voltage steps from –40 mV to –120 mV (10 mV increments) from a –40 mV holding potential. Current density was calculated by subtracting the instantaneous current (t = 0) from the steady-state current at the end of each voltage step.

#### Single-Cell/Nucleus RNA Sequencing

##### Isolation and processing of fetal tissue for single-nucleus RNA sequencing

To obtain AVN fetal tissue for single-nucleus RNA sequencing (snRNA-seq), we dissected the lower right atrium near the atrial–ventricular boundary from a fetal heart at 19 weeks of gestation. The tissue was immediately submerged in liquid nitrogen for rapid freezing and subsequently stored at −80 °C. For processing, the tissue was dissociated in a lysis buffer composed of 0.32 M sucrose (Sigma, S0389), 5 mM CaCl₂ (Fluka, 21114), 3 mM Mg(AC)₂ (Sigma, 228648), 20 mM Tris-HCl (ThermoFisher, 15567-027), 0.1% Triton X-100 (MilliporeSigma, X100), and 0.1 mM EDTA (ThermoFisher, AM9260G) prepared in DNase/RNase-free water (Qiagen, 129115). The tissue was mechanically homogenized with a douncer (Sigma, D9063) to yield single nuclei, which were then washed twice with PBS containing 1% BSA and 0.2 U/µl RNAseOUT (ThermoFisher, 10777019) and filtered through a 40 μm cell strainer. Finally, the nuclei were stained with DAPI (10 μg/ml) and sorted for DAPI^+^ single nuclei using a FACSAria (BD) at the SickKids-UHN Flow Cytometer Facility.

##### Isolation and Processing of differentiated AVNLPCs for single-cell RNA sequencing

To perform single-cell RNA sequencing on BMP2 AVNLPCs derived from the HES2 cell line, embryoid bodies (EBs) were dissociated into a single-cell suspension on day 25 of differentiation using the methods described above. The cells were then resuspended in 1X PBS containing 0.04% BSA and filtered through a 40 μm cell strainer.

##### Single-cell/nucleus RNA sequencing, raw data processing, quality control and clustering

Both single-cell and single-nuclus suspensions were prepared for sequencing using the 10x Genomics Chromium platform with the Single Cell 3’ v3 reagent kit. The resulting libraries were sequenced on an Illumina NovaSeq 6000, collecting over 45,000 reads per cell. Raw data were processed with the 10x Genomics CellRanger pipeline at UHN’s Princess Margaret Genomics Centre. For mapping, single-cell RNA-seq data were aligned to the human reference genome GRCh38, while single-nucleus RNA-seq data were aligned to GRCh38-premrna. All downstream analyses were performed in R using the Seurat toolkit. (version 5.0.1, https://satijalab.org/seurat/). First, Low-quality cells and potential doublets were filtered out. Specifically, cells with a library size below (10000 for hPSC and 3000 for fetal) or above 50,000 were excluded. Additionally, we removed damaged cells by applying mitochondrial gene expression thresholds (fetal snRNA-seq: 0.75% and hPSC scRNA-seq: 13.093%). The remaining data was then normalized using SCTransform. Next, we performed principal component analysis with RunPCA and selected the top 25 principal components for further clustering via the FindClusters function. The final cluster resolutions were set at 0.1 (full dataset) and 1.4 (cardiomyocyte subcluster) for the fetal snRNA-seq dataset and 0.1 (full dataset) and 0.5 (cardiomyocyte subcluster) for the hPSC scRNA-seq dataset.

##### Differential gene expression analysis

To identify differentially expressed genes (DEGs) within each cluster we employed the FindAllMarkers function with the following parameters: only.pos = TRUE, min.pct = 0.1, logfc.threshold = 0.25, and an adjusted p-value cutoff of <0.05. The most significant DEGs were selected by sorting based on adjusted p-value. To generate cardiomyocyte only datasets, we further subclustered cells from clusters that expressed *TNNT2* and defined the cardiomyocyte subtypes as detailed in the main text. To visualize gene expression in two dimensions Uniform Manifold Approximation and Projection (UMAP) was used for dimensionality reduction.

##### Gene Signature Scores

Gene signature scores were generated using UCell^94^ based on the top 50 positive DEGs from the comparison groups. When there were fewer than 50 positive DEGs, all were included. The distribution of these signature scores was visualized using UMAP projections. For the comparison to the published fetal spatial MERFISH dataset^59^ and adult AVN single nucleus datasets^95,96,63,97^ the top DEGs of each cell cluster of interest were identified in the supplementary data provided for each publication.

##### Gene ontology analysis

Database for Annotation, Visualization, and Integrated Discovery (DAVID) Bioinformatics Resources (version Dec. 2021, https://david.ncifcrf.gov/summary.jsp) was used to execute gene ontology analyses, focusing on biological processes (GOTERM_BP_all).

##### Spearman correlation analysis

To evaluate the correlation between selected fetal AVN cell types and hPSC-derived BMP2 AVNLPCs, we performed Spearman correlation analysis using the top 200 DEGs ranked by fold change. The analysis was conducted on specific cell clusters from each dataset. In the fetal dataset, these included the Core AVN, Atrial Transition Zone (ATZ), Lower Nodal Bundle (LNB), Atrial, Ventricular, Epithelial, and Fibroblast clusters. In the hPSC dataset, the analyzed clusters comprised the Core AVN, ATZ, LNB, Epithelial, Epicardial, and Fibroblast populations. Duplicate features and genes not detected in the hPSC dataset were excluded, and the remaining genes were used to calculate Spearman correlation scores between the fetal and hPSC clusters.

#### Engineered Heart Tissues

##### Cardiomyocyte purification for engineered heart tissues

To generate eAVNT and eVT from the HES3-Nkx2-5^egfp/w^ TBX3^tdtom/w^ hPSC line SIRPA^+^CD90^−^NKX2-5^+^TBX3^+^ AVNLPCs and SIRPA^+^CD90^−^NKX2-5^+^TBX3^−^ VLCMs were isolated by FACS respectively and directly used for tissue generation. To generate eAVNT and eVT from the ESI017-ASAP1 hPSC line cardiomyocytes were enriched up to 80% by MACS using the PSC-Derived Cardiomyocyte Isolation Kit (Miltenyi, 130-110-188). After the sort, the ESI017-ASAP1 AVNLPCs and VLCMs were cryopreserved as outlined above.

##### Cardiomyocyte thawing for engineered heart tissues

Cryopreserved vials of hPSC-derived cardiomyocytes were thawed in a 37°C water bath until a small ice pellet remained. Cells were gently resuspended in cardiac maintenance medium supplemented with 10 µg/mL DNase I (MilliporeSigma, 260913) and 10μM ROCK inhibitor Y27632 (Selleckchem, S1049), transferred to a 15 mL conical tube, and washed with the same medium. Cells were centrifuged at 1200 rpm for 5 minutes, and this process was repeated twice. The final pellet was resuspended in cardiac maintenance medium containing 10μM ROCK inhibitor and kept on ice until use.

##### Generation of engineered heart tissues

For the preparation of 3mm engineered heart tissues, PDMS moulds were prepared by securing 210 µm diameter PDMS rods at either end of a PDMS well (4mm long x 2mm wide x 0.6mm deep) to serve as anchors for the tissue. The mold dimensions were based on those of previously reported devices^95,96,63,97^. Cardiomyocytes were counted and seeded at a density of 160,500 cells per construct, along with commercially available cardiac fibroblasts (Lonza, CC-2904) at a density of 53,500 cells per construct. The cells were mixed and embedded in fibrin gels composed of fibrinogen (5 mg/mL; Sigma cat# F8630), aprotinin (0.00825 mg/mL; BioShop cat# APR600), thrombin (2 U/mL; Sigma-Aldrich cat# T7513), growth factor-reduced Matrigel (1 mg/mL), and Y-27632 (10 nm, STEMCELL cat# 72304), in DMEM. The cell suspension was seeded into the molds by pipetting with cold tips to prevent premature fibrin polymerization. The gels were incubated at 37°C for 1 hour to allow for polymerization, after which cardiac maintenance media (see above) was added. The tissues were cultured for 7 days to enable compaction, with the medium replaced every 2–3 days. Five days after seeding, the tissues were carefully detached from the walls of the molds using a 20-gauge needle to promote further compaction. Following an additional 5 days in culture (10 days total), the tissues were optically mapped.

##### Optical mapping of engineered heart tissues

Engineered heart tissues were mapped in 20% StemPro-34 diluted with IMDM supplemented with the above-described culture reagents. Tissues made with cells differentiated from the HES3-Nkx2-5^egfp/w^ TBX3^tdtom/w^ hPSC line were stained at 37°C, 5% CO_2_ using the Fluo4 Membrane Potential Kit (ThermoFisher, F14201) according to the manufacturer’s instructions. Fluorescence was recorded using a high-speed complementary metal-oxide semiconductor (CMOS) camera (Ultima-L, Scimedia) equipped with a 1cm sensor with a 100 × 100-pixel resolution, mounted on an MVX-10 Olympus fluorescence microscope with a 0.63-6.3X objective lens. Optical zoom was set at 4X, providing a 25um/pixel spatial resolution for imaging. Recordings of 2-8 seconds were acquired at 500 or 1000 frames/s using the BrainVision software (Scimedia). The calcium indicator Fluo4 and the fluorescent voltage sensor ASAP1 were excited by a mercury arc source (X-Cite Exacte) with a 480nm filter (Olympus U-MWIG2 filter cube). To detect fluorescence a 488nm long pass filter was used. A heated plate (MATS-U55S, Olympus) was used to maintain the media/buffer at 37 °C.

The tissues were electrically point-stimulated using a bipolar electrode made of two fine silver wires supported by a syringe mounted on a micromanipulator (World Precision Instruments) and connected to a Grass S88X stimulator. Tissues were paced at 2.5 Hz to determine conduction velocities. To test for the ability of the tissues to block the conduction of fast rates they were stimulated for 2 seconds at each frequency ranging from 2.5 Hz to 10 Hz. Decremental conduction properties were assessed using the following S1-S2 stimulation protocol. 8 stimuli (S1) at a fixed frequency of 2.5 Hz were applied to the tissues followed by an extra stimulus (S2) with increased frequency ranging from 3 Hz to 10 Hz (Figure S6B). The time (also referred to as latency) from the S2 stimulus to the activation of the voltage/calcium transient (R2) was determined for each S2 frequency to assess for decremental conduction (Figure S6C).

Optical action potential characteristics (frequency, maximal upstroke velocity, and action potential duration) were analyzed using the Electromap Matlab software^98^. Activation maps were also created using Electromap. Conduction properties (conduction velocity, maximum capture rate, decremental conduction and the instance of the Wenckebach phenomenon) and diastolic depolarization were determined using the BrainVision software (Scimedia).

#### Guinea Pig Experiments

##### Guinea pig cryoinjury

Guinea pig experiments, including cardiac cryoinjury, cell injection, and ex vivo imaging, were performed as described in previous studies^56,68,69^. Briefly, animals were initially anesthetized with ketamine and xylazine, then maintained under anesthesia with 1.5% isoflurane and oxygen via intubation and mechanical ventilation (60 breaths per minute, tidal volume 5 mL/kg). A cryoinjury was induced by performing a left thoracotomy to expose the heart, followed by the application of a 5-mm diameter liquid nitrogen-cooled aluminum cryoprobe to the anterolateral left ventricle (LV) four times, each for 30 seconds.

##### Cardiomyocyte thawing for in vivo transplantation

Cryopreserved cardiomyocytes were thawed and prepared following previously established protocols to maximize survival during in vivo transplantation^56,68,69^. Briefly, cryopreserved vials of hPSC-derived cardiomyocytes were retrieved from a -180°C freezer and rapidly thawed in a 37°C water bath until a small ice pellet remained. Cells were gently resuspended and washed in pre-cooled 20% StemPro medium diluted in IMDM without Phenol Red (Thermo Fisher, 21056023), supplemented with 200 nM cyclosporine A/Sandimmune IV (Novartis, 00593257), 50 µM Pinacidil (Sigma, P154-100MG), and 10 µg/mL DNase I (MilliporeSigma, 260913). Cells were then pelleted by centrifugation at 1200 rpm for 10 minutes. The pellet was resuspended in pre-chilled IMDM without Phenol Red, supplemented with 200 nM cyclosporine A/Sandimmune IV, 50 µM Pinacidil, and 10 µg/mL DNase I, followed by centrifugation at 300 g for 10 minutes. This wash step was repeated twice. Finally, cells were resuspended in 100% cold Matrigel without Phenol Red (FisherScientific, CB40230C) supplemented with 200 nM cyclosporine A/Sandimmune IV and 50 µM Pinacidil to the target total injection volume of 250ul. All pipetting steps were performed using cold, wide-bore pipette tips to minimize mechanical stress. Cell suspensions were kept on ice and sealed with parafilm to prevent contamination until transplantation.

##### Guinea pig cell injection

A second thoracotomy was conducted 10-14 days post-cryoinjury, during which 50 -100 million hPSC-derived AVNLPCs or VLCMs were injected aiming to achieve grafts spanning from the scar into the healthy myocardium. Specifically, 2-3× 100uL injections into the scar and border zone of the scar were performed. To prevent graft rejection, animals were treated with pharmacological immunosuppression starting 2 days before cell injection and continuing until the endpoint of the experiment. The immunosuppression regiment consisted of cyclosporine A/Sandimmune IV (15 mg/kg/day subcutaneously until day 7 post-transplant, followed by 7.5 mg/kg/day until day 15 post-transplant, and 5 mg/kg/day until the endpoint, Novartis, 00593257) and methylprednisolone (2 mg/kg/day intraperitoneally, Pfizer, DIN02367971).

##### Guinea pig ex vivo optical mapping

At 2-4 weeks post-transplantation, hearts were excised and mounted onto a Langendorff perfusion system. For ex vivo maintenance, the hearts were perfused with Working Heart Buffer containing 118 mM NaCl, 1.8 mM CaCl₂, 4.7 mM KCl, 1.2 mM MgSO₄, 1.2 mM KH₂PO₄, 25 mM NaHCO₃, 11 mM D-glucose, and 1 mM sodium pyruvate (Key Resources ***Table S8***). The buffer was bubbled with 95% O₂/5% CO₂, adjusted to pH 7.35, and maintained at 37°C. To minimize motion artifact during ex vivo imaging, the buffer was supplemented with 10 µM blebbistatin (Cayman Chemical, 13186-50). A PowerLab 8/35 Data Acquisition System (AD Instruments) outfitted with a bioamplifier (Model ML136, AD Instruments), was used to record a pseudo-ECG via the Labchart software. The voltage-sensitive dye RH237 (Ex/Em: 532/700nm) was used to detect optical action potentials of the host. For the simultaneous recording of ASAP1 (hPSC-derived cardiomyocyte graft) and RH237 (guinea pig heart) fluorescent signals, a previously reported dual camera imaging system was used. This comprised of a collimating lens (25 mm, Navitar, N.A. 0.05, W.D. 10mm) outfitted with a dichroic system (Photometrics DC2 dual-channel splitter) and two high-speed, high-sensitivity EMCCD cameras (Evolve128, Photometrics). The cameras were operated by the Metamorph software at 500 fps. Two LED spotlights (Mightex PLS-0470-030-15-S LEDs with Chroma ET470/40x filters) were used to excite the fluorescent signals. Prior to detection, the ASAP1 and RH237 fluorescent signals were separated by a dichroic mirror (643 nm) and filtered (530 ±20 nm and >705 nm, respectively).

The hearts were mapped at sinus rhythm and after the induction of an SAN/AVN block, which was triggered by adding methacholine (10 mg/ml, Sigma, A2251-25G) to the perfusate. The hearts or cellular grafts were electrically point-stimulated using a bipolar electrode made of two fine silver wires (AWG#32) supported by a syringe mounted on a magnetic stand with a ball joint (World Precision Instruments) driven by the PowerLab system. Hearts/grafts were paced at 2.0 Hz to determine conduction velocities within the grafts. To test for the ability of grafts to block the conduction of fast rates heart/grafts were paced with increasing frequencies ranging from 1 Hz to 8 Hz in 0.5Hz increments.

Raw data from the ASAP1, RH237, and ECG channels were processed using previously described Matlab scripts to generate ***Figure 7D-7E, 7L and 7M***^56^. Optical action potential characteristics, including frequency, maximum upstroke velocity, and action potential duration, as well as conduction velocity, were analyzed with Electromap software^98^. Activation maps were also created using Electromap. Maximum capture rates and optical action potential analysis for diastolic depolarization were performed using Metamorph software.

### STATISTICAL ANALYSIS

Data are presented as mean ± standard error of the mean (SEM). For all In vitro differentiations indicated samples size (n) represent biological replicates (independent differentiation experiments). For in vivo animal studies, experimental group size (n = animal numbers) was estimated based on power analyses using population variance in previous work. Animals were selected at random by a technician blinded to experimental treatment. Statistical analysis was performed using GraphPad Prism version 10.4.1 for macOS (www.graphpad.com). Datasets were assessed for normality using the Shapiro-Wilk test (α = 0.05). For normally distributed data, the following statistical tests were employed: a two-tailed Student’s t-test for comparisons between two conditions, and one-way ANOVA with Bonferroni post hoc test for comparisons among multiple conditions. For datasets that did not pass the normality test, nonparametric statistical tests were used. Results were considered statistically significant at p < 0.05 (*), p < 0.01 (**) and p < 0.001 (***). All sample sizes, statistical methods and statistical parameters, including p-values, are detailed in the respective figures and figure legends. Due to the nature of the study, the experiments were not randomized and the investigators were not blinded to allocation during experiments and outcome assessment. Except for the animal experiments, no statistical methods were used to pre-determine sample sizes.

### DATA AVAILABILITY

The data supporting the findings of this study are available within the article and the supplemental information files. Additional information is available from the corresponding author upon request.

### CODE AVAILABILITY

This study did not involve the development of original code. All software packages utilized are described and cited in the Methods section.

### FIGURE PREPARATION

Adobe Illustrator was used to prepare figures. Biorender was used to generate schematics.

## ACKNOWLEDGEMENTS

We would like to thank all members of the Protze laboratory for experimental advice and critical comments on the manuscript. We thank Prof. A. Elefanty and Prof. E. Stanley (Monash University, Victoria, AU) for providing the HES3 NKX2-5^egfp/w^ reporter line. We thank Dr. Faisal Alibhai (University Health Network, Toronto, CA) for insightful feedback and guidance on the optical mapping analysis. We would also like to thank the core facilities that supported this project including: the SickKids-UHN Flow Cytometry Facility for assistance with cell sorting, the Advanced Optical Microscopy Facility at UHN for assistance with confocal microscopy, the Princess Margaret Genomics Centre at UHN for assistance with sc/snRNA-seq and data processing and the Animal Resource Centre at UHN for veterinary assistance with animal experiments. We thank the donors, the Research Centre for Women’s and Infants’ Health BioBank Program, the Lunenfeld-Tannenbaum Research Institute, and the Department of Obstetrics & Gynecology of Sinai Health System for the human specimens used in this study. This work was supported by grants from the Canadian Institutes of Health Research (CIHR PJT 169090, S.P.), the Canadian Foundation for Innovation (John R. Evans Leaders Fund, 38651, S.P.), the Government of Canada’s New Frontiers in Research Fund (NRFT-2022-00447, M.A.L.)”, the Canada Research Chairs Program (CRC-2020-00245, M.A.L.), and funding from BlueRock Therapeutics LP (S.P.). M.L. was supported by a Canadian Institutes of Health Research Masters Fellowships and Ontario Graduate Scholarships. A.A.L., and A.X. were supported by Ontario Graduate Scholarships, B.M.M. and M.L.C. were supported by Canadian Institutes of Health Research Doctoral Fellowships.

## AUTHOR CONTRIBUTIONS

M.L. designed the project, performed experiments, analyzed data, and wrote the manuscript. M.L., M.K. and R.S. performed and analyzed tissue culture and flow cytometry experiments. O.Mo., O.Ma., K.S., and S.N.V. designed and generated the engineered heart tissues. S.P. designed, performed, and analyzed the patch clamp experiments. M.L., S.P., S.M., and K.N. designed, performed and analyzed the optical mapping experiments of the engineered heart tissues. M.L., B.M.M., S.P., W.D., and M.A.L. designed, performed and analyzed the ex vivo optical mapping experiments of the guinea pig hearts. A.X. assisted with analysis of optical mapping data and optical action potential parameters. M.E. and B.Q. assisted with the guinea pig surgeries and cell injections. M.L.C. assisted with immunohistochemistry experiments and imaging of guinea pig heart sections. S.K. designed and generated the NKX2-5^eGFP/wt^ TBX3^tdTomato/wt^ hPSC reporter line. A.M., assisted with the dissection of fetal heart tissues. A.A.L., M.L., and S.P. generated and analyzed the sc/snRNA-seq data. S.P. designed the project and wrote the manuscript.

## MATERIALS & CORRESPONDENCE

Any request for material should be addressed to S.P.

## Supplemental Figure Legends

**Supplementary Figure 1:**
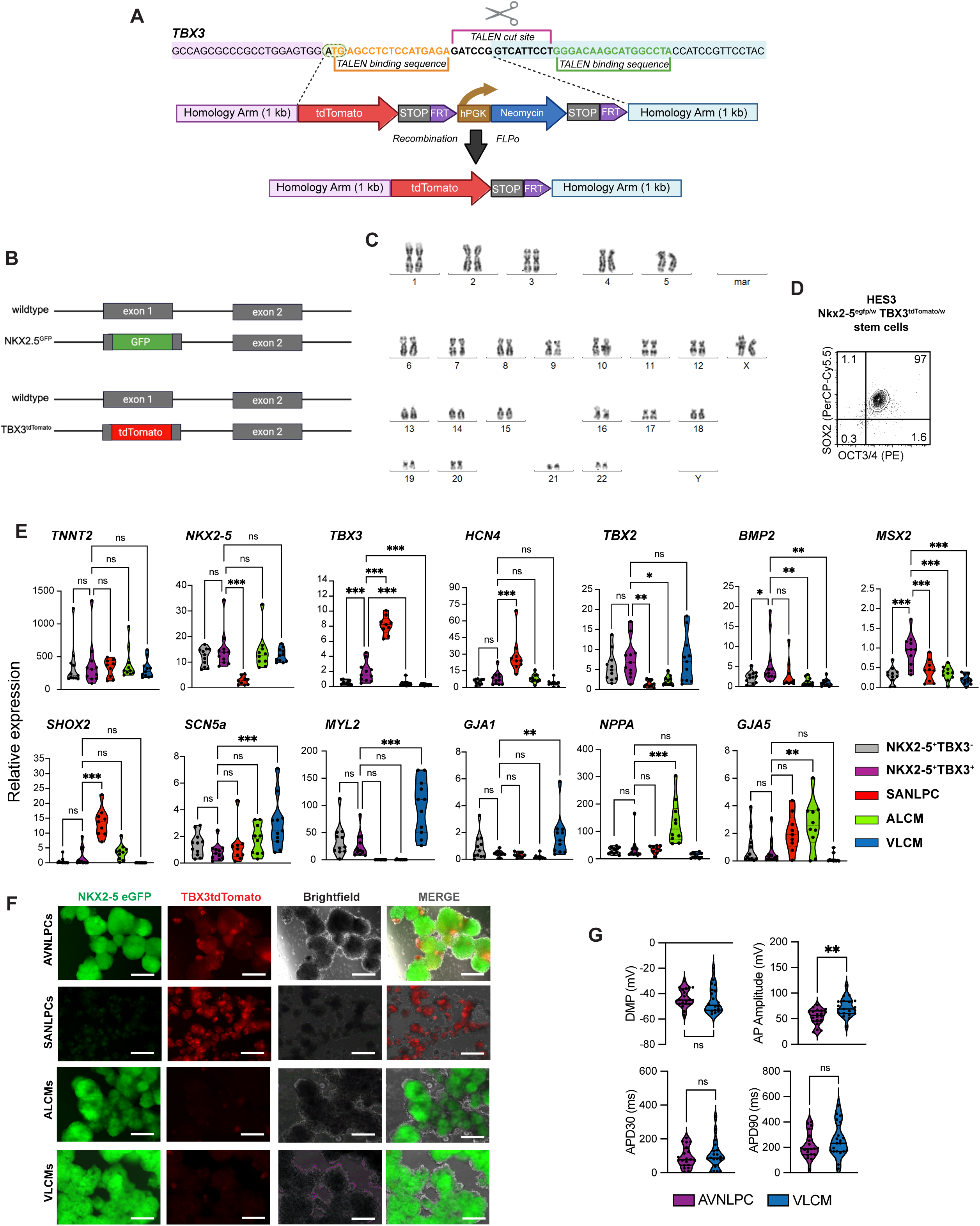
NKX2-5⁺ TBX3⁺ cells exhibit the molecular signature of and functionally resemble the endogenous AVN. Related to Figure 1. **A.** Schematic of TALEN-based insertion of tdTomato reporter into the 1^st^ exon of one allele of TBX3. TBX3 genomic sequence with TALEN binding site (top). Targeting vector (middle). Targeted TBX3 allele after removal of Neomycin selection cassette (bottom). **B**. Schematic of final double knock-in HES3-Nkx2-5^egfp/w^ TBX3^tdTomato/w^ hPSC reporter line. **C.** Karyogram of HES3-Nkx2-5^egfp/w^ TBX3^tdTomato/w^ hPSC line (46, XX). **D.** Flow cytometric analysis of OCT3/ 4 and SOX2 expression in the HES3-Nkx2-5^egfp/w^ TBX3^tdtom/w^ hPSC line. **E.** Violin plots of the RT-qPCR analysis presented as heat map in Fig. 1H (n = 7-8). Values represent expression relative to the housekeeping gene TBP. **F.** Representative fluorescent and brightfield images for the expression of NKX2-5:GFP and TBX3:tdTomato of day 20 NKX2-5^+^TBX3^+^ AVNLPCs, SANLPCs, ALCMs, and VLCMs. Scale bars, 250 μM. **G.** Quantification of indicated action potential parameters. (NKX2-5^+^TBX3^+^ AVNLPC: n = 20, VLCM: n = 20). Error bars represent SEM. Statistical tests: One-way ANOVA, Bonferroni’s post hoc test (E); Two-sided t-test (G); Datasets that failed the normality test (G: DMP, APD30) were analyzed using Mann-Whitney test: *P < 0.05, **P < 0.01, ***P < 0.001. AP, action potential; APA, AP amplitude, APD30/90, AP duration at 30%/90% of repolarization; DMP, Diastolic membrane potential.

**Supplementary Figure 2:**
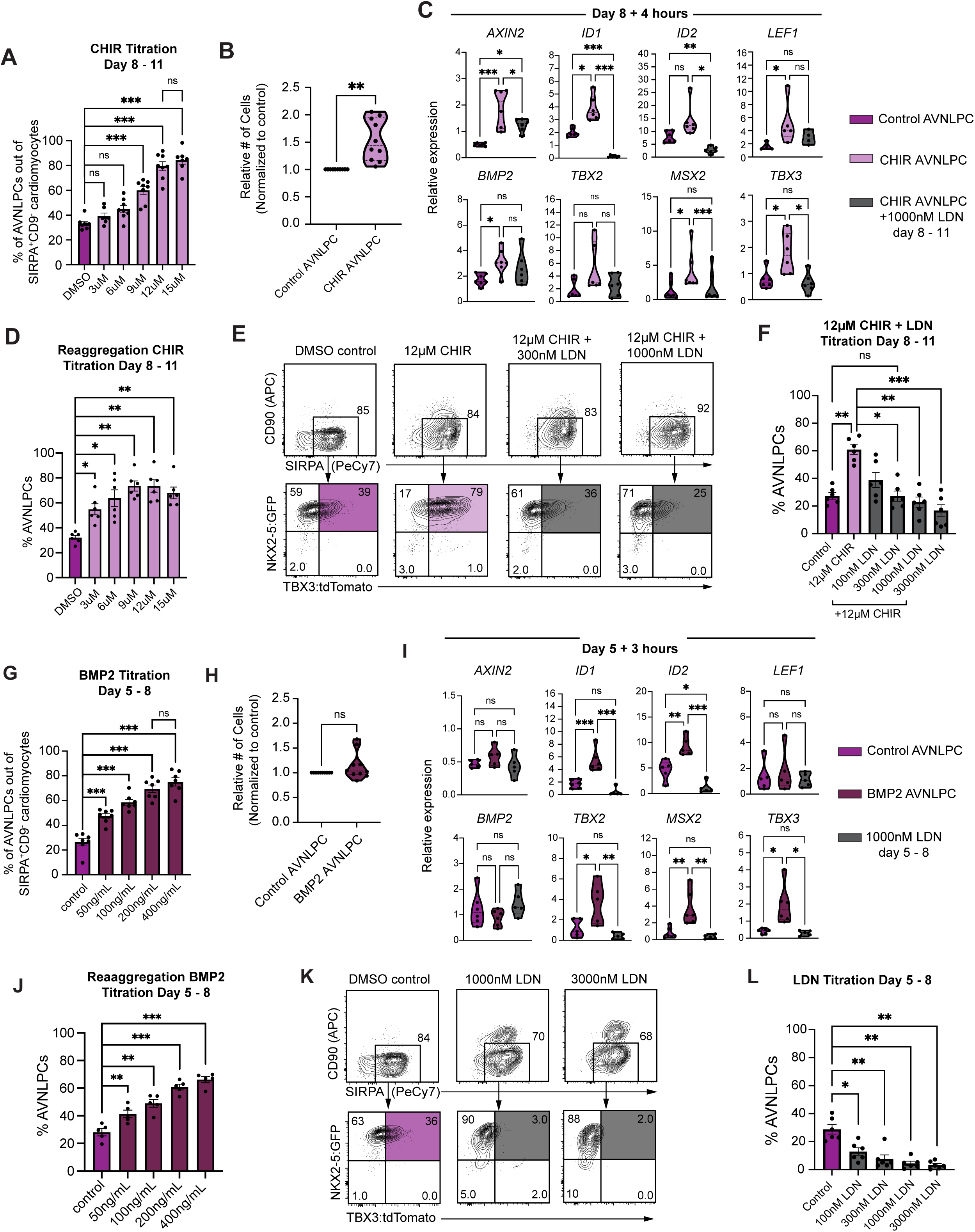
Modulating AVN-specific developmental signaling pathways. Related to Figure 2. **A.** Bar graph summarizing the proportion of NKX2-5^+^TBX3^+^ cells gated out of SIPRA^+^CD90^−^ cardiomyocytes after treatment with the indicated amounts of CHIR at days 8-11 as shown in Fig. 2D (n = 5). **B.** Violin plot showing the relative number of cells per well of CHIR AVNLPCs on Day 20 of differentiation normalized to Control AVNLPCs (n=10). **C.** RT-qPCR analysis of EBs isolated 4 hours after treatment with 12 μM CHIR (pink) and 12 μM CHIR + 300nM LDN (grey) at day 8 of differentiation for the expression of the indicated genes. Control AVNLPCs (purple) were included as expression reference (n = 6-7). Values represent expression relative to the housekeeping gene TBP. **D.** Proportion of NKX2-5^+^TBX3^+^ cardiomyocytes out of total cells after treatment with the indicated amounts of CHIR at days 8-11 following a reaggregation step. **E.** Representative day 20 flow cytometric analysis of NKX2-5:GFP and TBX3:tdTomato expression in SIPRA^+^CD90^−^ cardiomyocytes after treatment with 12 μM CHIR and the indicated amount of LDN days 8-11 of differentiation in comparison to the untreated control. **F.** Bar graph summarizing the proportion of NKX2-5^+^TBX3^+^ cardiomyocytes out of total cells shown in E. **G.** Bar graph summarizing the proportion of NKX2-5^+^TBX3^+^ cells gated out of SIPRA^+^CD90^−^ cardiomyocytes after treatment with the indicated amounts of BMP2 at days 5-8 of differentiation as shown in Fig. 2H (n = 7). **H.** Violin plot showing the relative number of cells per well of BMP2 AVNLPCs on Day 20 of differentiation normalized to Control AVNLPCs (n = 10). **I.** RT-qPCR analysis of EBs isolated 3 hours after treatment with 200 ng/mL BMP2 (maroon) and 1000nM LDN (grey) at day 5 of differentiation for the expression of the indicated genes. Control AVNLPCs (purple) were included as expression reference (n = 6-7). Values represent expression relative to the housekeeping gene TBP. **J**. Proportion of NKX2-5^+^TBX3^+^ cardiomyocytes out of total cells after treatment with the indicated amounts of BMP2 at days 5-8 following a reaggregation step (n = 5). **K.** Representative day 20 flow cytometric analysis of NKX2-5:GFP and TBX3:tdTomato expression in SIPRA^+^CD90^−^ cardiomyocytes after treatment with the indicated amounts of LDN at days 5-8 of differentiation. **L.** Bar graph summarizing the proportion of NKX2-5^+^TBX3^+^ cardiomyocytes out of total cells shown in K. Error bars represent SEM. Statistical tests: One-way ANOVA, Bonferroni’s post hoc test (A, C, D, F, G, I, J, L); unpaired two-sided t-test (B, H): *P < 0.05, **P < 0.01, ***P < 0.001. CHIR, CHIR99021; LDN, LDN-193189.

**Supplementary Figure 3:**
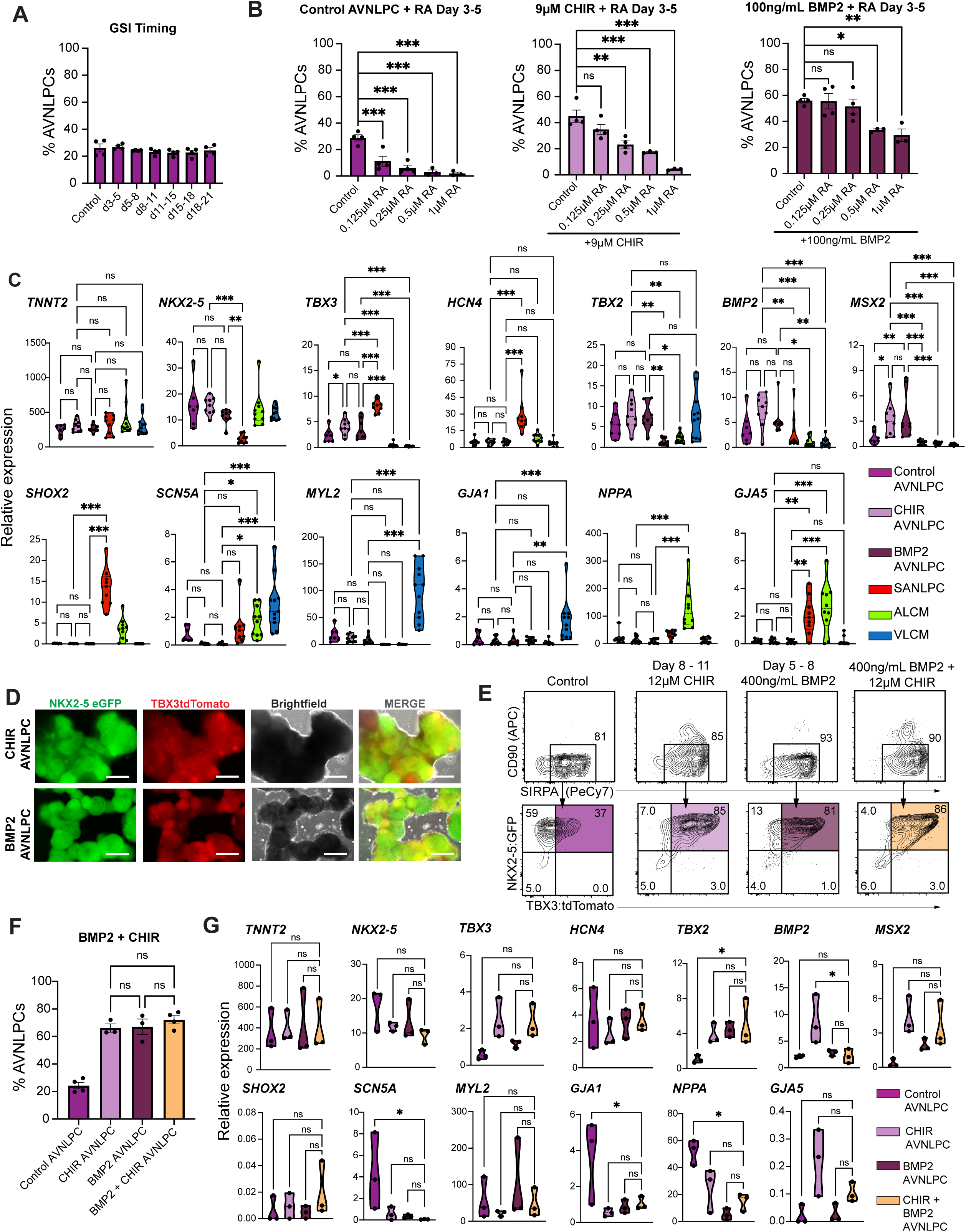
NOTCH inhibition, RA treatment and combined BMP2 + CHIR treatment do not further increase AVNLPC content. Related to Figure 2. **A.** Bar graph summarizing the proportion of NKX2-5^+^TBX3^+^ cardiomyocytes out of total cells after treatment with 10 μM GSI during the indicated timepoints in comparison to the untreated control. (n = 4). **B.** Bar graphs summarizing the proportion of NKX2-5^+^TBX3^+^ cardiomyocytes out of total cells after treatment with the indicated amounts of Retinoic Acid at days 3-5 of differentiation in control AVNLPCs (left), AVNLPCs treated with 9 μM CHIR at days 8-11 (middle) and AVNLPCs treated with 100 ng/mL of BMP2 at days 5-8 (right) (n = 3-4). **C.** Violin plots of the RT-qPCR analysis presented as heat map in Fig. 2K (n = 7-8). Values represent expression relative to the housekeeping gene TBP. **D.** Representative fluorescent and brightfield images for the expression of NKX2-5:GFP and TBX3:tdTomato of day 20 CHIR AVNLPCs and BMP2 AVNLPCs. Scale bars, 250 μM. **E.** Representative day 20 flow cytometric analysis of NKX2-5:GFP and TBX3:tdTomato expression in SIPRA^+^CD90^−^ cardiomyocytes after treatment with 12 μM CHIR days 8-11, 400 ng/mL BMP2 days 5-8 or both 400ng/mL BMP2 days 5-8 and 12 μM CHIR days 8-11 (BMP2 + CHIR AVNLPCs) in comparison to untreated control. **F.** Bar graph summarizing the proportion of NKX2-5^+^TBX3^+^ cardiomyocytes out of total cells shown in E (n = 3-4). **G.** RT-qPCR analysis of NKX2-5^+^TBX3^+^SIRPA^+^CD90^−^ sorted BMP2 + CHIR AVNLPCs (yellow) for the expression of the indicated genes. NKX2-5^+^TBX3^+^SIRPA^+^CD90^−^ sorted Control AVNLPCs (purple), CHIR AVNLPCs (pink) and BMP2 AVNLPCs (maroon) are included as expression reference. All populations were isolated at day 20 of differentiation (n = 3). Values represent expression relative to the housekeeping gene TBP. Error bars represent SEM. Statistical tests: One-way ANOVA, Bonferroni’s post hoc test (B, C, F, G): *P < 0.05, **P < 0.01, ***P < 0.001. GSI, Gamma Secretase Inhibitor; RA, Retinoic Acid; CHIR, CHIR99021

**Supplementary Figure 4:**
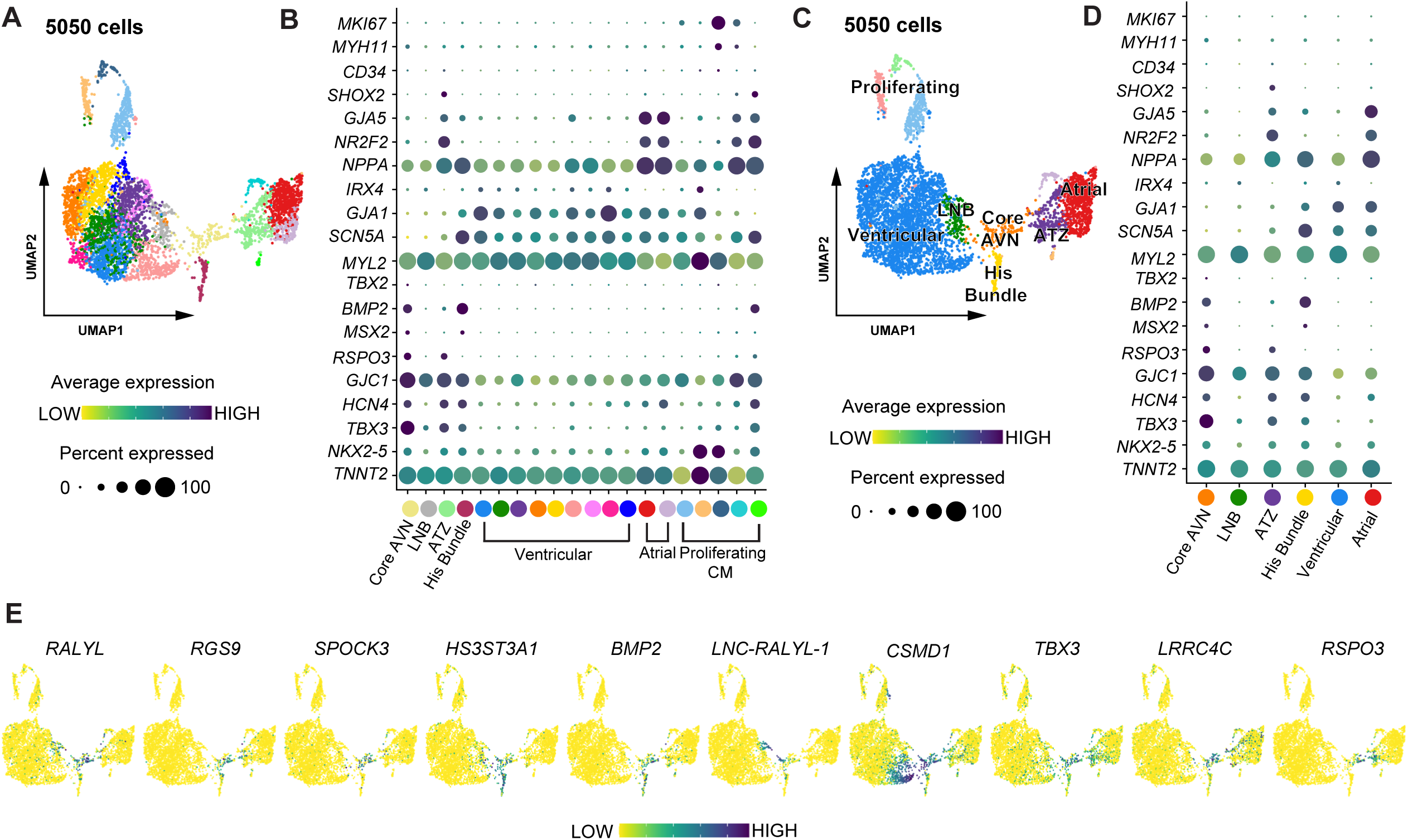
Subclustering of fetal AVN cardiomyocytes. Related to Figure 4. **A.** UMAP of Subclustered *TNNT2*^+^ cardiomyocytes showing 20 cell clusters. **B.** Dot plot depicting the expression of the indicated genes within the subclustered cardiomyocytes shown in A. **C.** UMAP showing merged subclusters based on the similarity in marker gene expression. **D.** Dot plot depicting the expression of the indicated genes within the merged cardiomyocyte subclusters shown in C. **E.** UMAPs showing the expression of the top 10 DEGs of the fetal AVN Core AVN cluster.

**Supplementary Figure 5:**
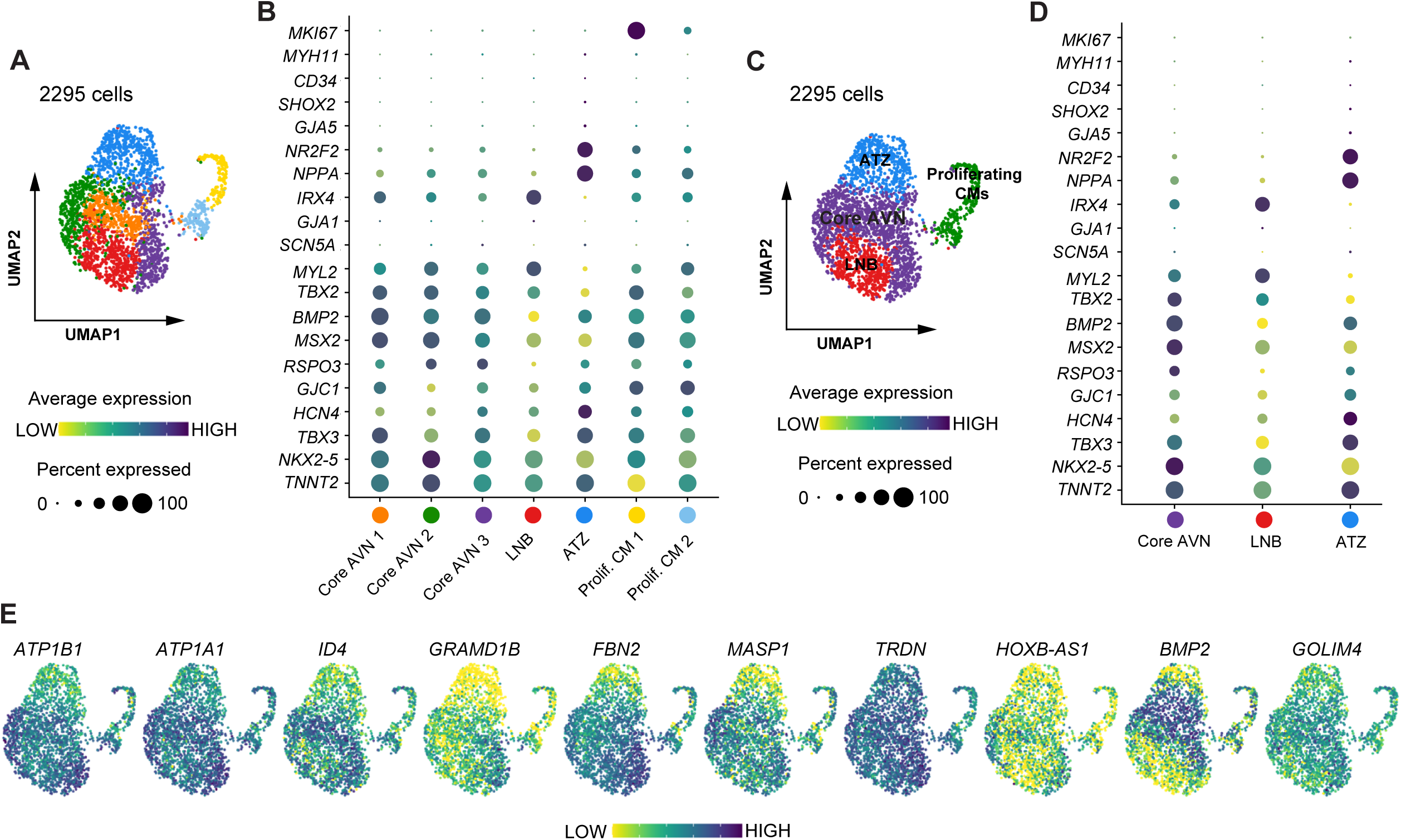
Subclustering of hPSC-derived AVNLPCs. Related to Figure 5. **A.** UMAP of subclustered *TNNT2***^+^** cardiomyocytes showing 7 distinct sub-clusters. **B.** Dot plot depicting the expression of the indicated genes within the subclustered cardiomyocytes shown in A. **C.** UMAP showing merged subclusters based on the similarity in marker gene expression. **D.** Dot plot depicting the expression of the indicated genes within the merged cardiomyocyte subclusters shown in C. **E.** UMAPs showing the expression of the top 10 DEGs of the hPSC-derived AVNLPC Core AVN cluster.

**Supplementary Figure 6:**
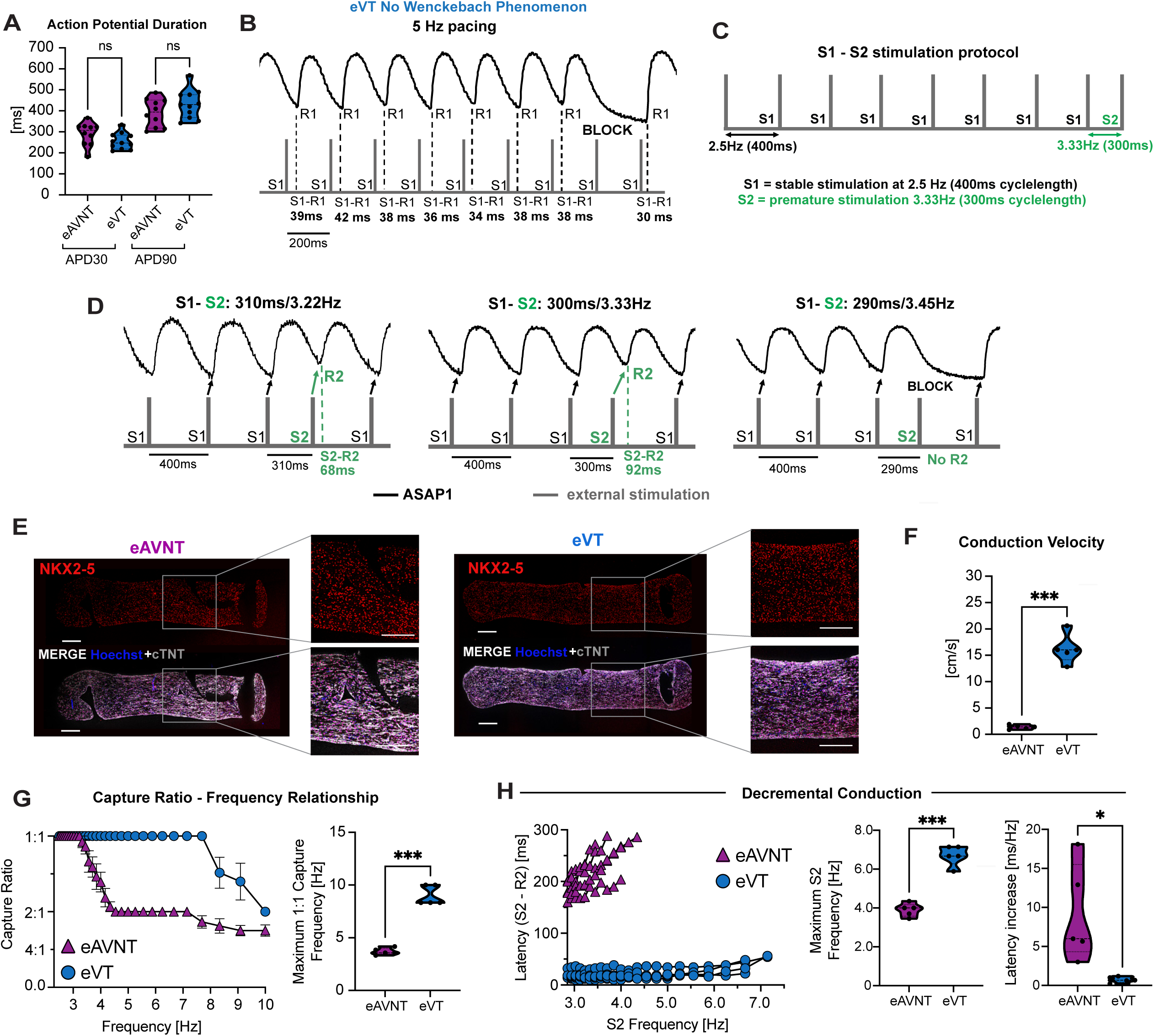
Engineered AVN tissues exhibit key AVN-like functional properties. Related to Figure 6. **A.** Quantification of indicated optical action potential duration at 30% and 90% of repolarization (eAVNTs: n= 2-3, eVTs: n=2-3, from four independent differentiations each). **B.** Representative optical action potential trace of an eVT not displaying the Wenckeback Phenomenon leading up to a blocked beat when paced at 5Hz. S1 (grey) denotes the applied stimulus via the pacing electrode, and R1 represents the tissue’s action potential response to S1. S1-R1 quantifies the impulse conduction time. **C.** Example of S1-S2 stimulation protocol to assess decremental conduction properties. A stable S1 stimulation is applied at 400ms cycle length (2.5Hz) at the proximal end of the tissue. Every 9 beats an extra S2 stimulation was applied with decreasing S1-S2 cycle length ranging from 350 to 100ms (2.86Hz to 10Hz). **D.** Representative ASAP1 voltage transients in an eAVNT at different S1-S2 cycle length. Arrows indicate ASAP1 transients (black) evoked by respective stimulation (grey). To quantify the conduction delay (latency), the time from S2 stimulation to ASAP1 transient activation (R2) was analyzed at the distal end of the tissue. **E.** Immunofluorescent staining of NKX2-5 in eAVNTs and eVTs counterstained with cTNT to identify cardiomyocytes and Hoechst to visualize all cells. Scale bars, 250 μm. **F-H.** Conduction properties recorded in eAVNTs and eVTs from the HES3-Nkx2-5^egfp/w^ TBX3^tdtom/w^ hPSC line (eAVNTs: n = 1-3, eVTs n = 1-3 from 3-5 independent differentiations each). **F.** Quantification of conduction velocities. **G.** Graph summarizing the capture ratios at increasing pacing frequencies (left). Quantification of the frequency at which 1:1 capture is lost (right). **H.** Assessment for decremental conduction properties. Graph displaying the response of the impulse conducting time (Latency, S2-R2) to increasing frequency of the extra stimulus (S1-S2) (left). Quantification of the frequency of the extra stimulus (S2) at which capture is lost (middle). Quantification of the slope of the latency increase (right). Statistical tests: unpaired two-sided t-test. APD30/90, action potential duration at 30/90% of repolarization.

**Supplementary Figure 7:**
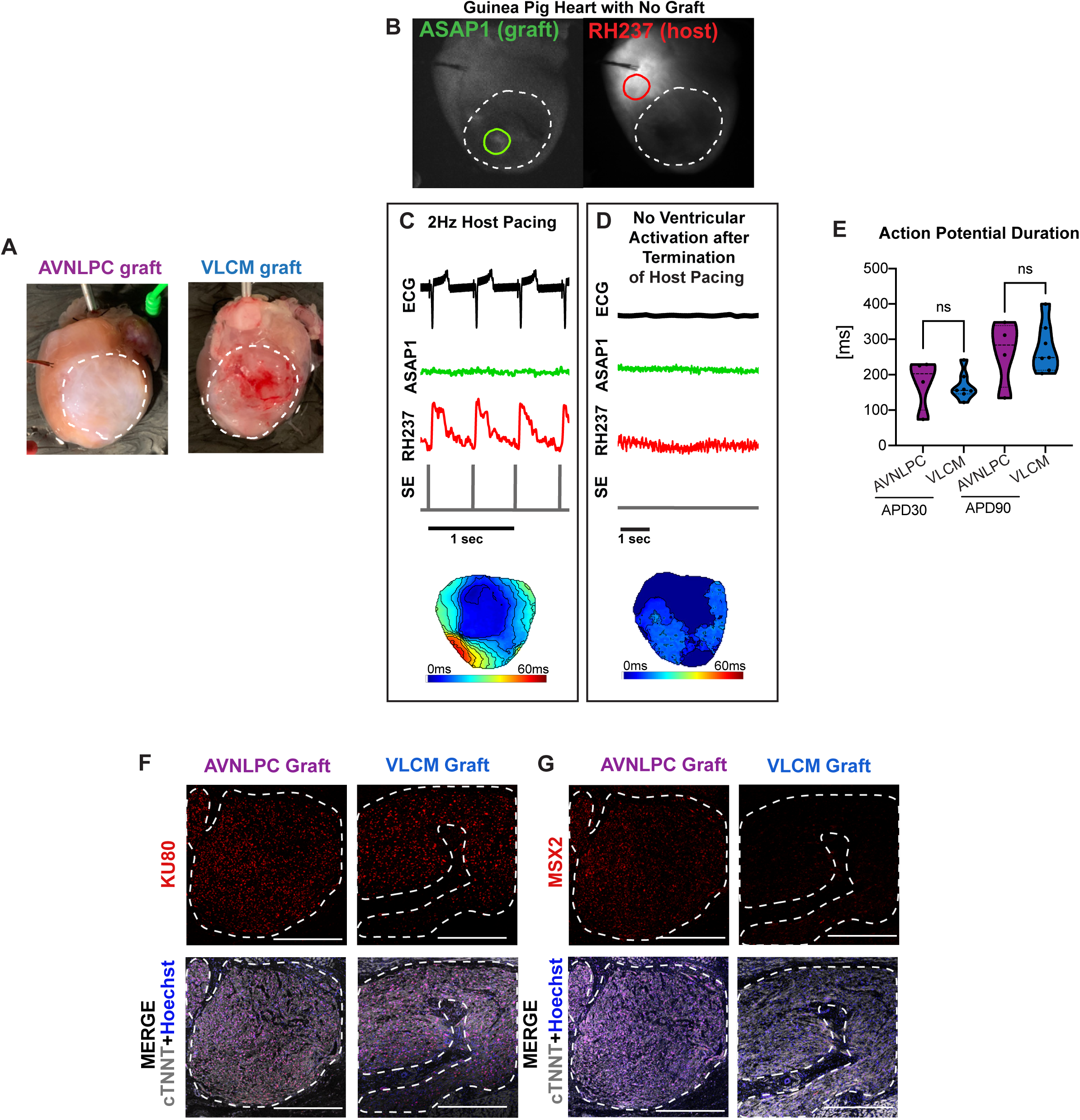
AVNLPCs retain their AVN-like functional properties in vivo. Related to Figure 7. **A.** Brightfield images of hearts with cryoinjury-induced scars. White dotted lines outline the scar area. **B.** Representative images of hearts without an ASAP1 hPSC-derived graft acquired in the ASAP1 (green) and RH237 (red) channels. The white dotted line indicates the scar area. Optical action potentials (oAP) of the ASAP1 signal are recorded in the region of interest (ROI) marked by the green solid line, while host oAPs are recorded in the ROI marked by the red solid line. **C, D.** Representative host ECG traces together with oAPs recorded in the ASAP1 (green) and RH237 (red, host) channels and the signal form the stimulation electrode (grey) showing electrode pacing at 2Hz, C. When external pacing was terminated (flat grey line) no electrical signal was detected in the ventricle due to methacholine effectively blocking conduction of endogenous pacing via the AVN, D. The bottom of each panel shows the voltage activation map of the heart derived from the RH237 signal indicating the origin of pacing. **E.** Quantification of optical action potential duration (APD) at 30% and 90% of repolarization in AVNLPC and VLCM grafts during spontaneous graft pacing (AVNLPC grafts: n = 4, VLCM grafts: n = 7). **F.** Immunofluorescent staining of KU80 in AVNLPC and VLCM grafts, counterstained with cTNT to identify cardiomyocytes and Hoechst to visualize all cells. Scale bars, 400 μm. **G.** Immunofluorescent staining of MSX2 in AVNLPC and VLCM grafts, counterstained with cTNT to identify cardiomyocytes and Hoechst to visualize all cells. Scale bars, 400 μm. Error bars represent SEM. APD30/90, action potential duration at 30/90% of repolarization.

